# The hidden diversity: Replication-strand asymmetry shapes short-term nucleotide and structural variation in an RNA virus

**DOI:** 10.64898/2026.09.04.749430

**Authors:** María J. Olmo-Uceda, Dominik Herek, Victoria G. Castiglioni, Santiago F. Elena

## Abstract

**Background:** Orsay virus (OrV), a positive-sense RNA virus infecting *Caenorhabditis elegans*, establishes latent infections with minimal impact on host fitness. Using strand-specific RNA-seq across twelve time points spanning larval development, we characterized OrV replication dynamics and short-term evolutionary patterns.

**Results:** Viral accumulation followed four distinct phases and exhibited strong strand asymmetry, consistent with a predominance of the stamping-machine replication mechanism. Diversity analyses revealed that negative-strand genomes harbor more mutations than positive strands, with hotspots concentrated in the replicase C-terminal tail and structural protein regions. Many negative-strand variants failed to propagate to positive strands, suggesting within-cell constraints or purifying selection at the RNA level. We also report the first evidence of non-standard viral genomes in OrV, dominated by deletions in RNA2, including a recurrent large deletion that may function as a subgenomic RNA encoding a shorter version of the δ protein.

**Conclusions:** These findings highlight replication asymmetry, strand-specific mutational filtering, and pervasive non-standard genomes as key features shaping OrV evolution during a primary infection.

## BACKGROUND

Viral populations within an infected host are highly dynamic, shaped by rapid replication, high mutation rates and intense competition among closely related variants (Cuevas et al. 2015; Nelson and Hughes 2015; Raghwani et al. 2019; Lauring 2020; Du et al. 2022). These processes generate mutant swarms whose composition reflects the balance between stochastic mutational input and selective forces operating at cellular and tissue scales, and they can now be quantified with precision thanks to advances in deep sequencing and analysis of within-host variation (Andino and Domingo 2015; Lauring 2020; Sardanyés et al. 2024). Conceptual and practical frameworks for measuring such diversity (*e.g*., allele frequency-based indices, entropy measures, and phylogenetic summaries) have coalesced over the last decade, enabling rigorous, comparable assessments across viruses and datasets (Lauring 2020; Fuhrman et al. 2021). These approaches have shown that within-host genetic diversity is not merely descriptive: it encodes how replication, mutation, drift, selection, and host factors jointly shape short-term viral evolution (Nelson and Hughes 2015; Lauring 2020).

A central insight from within-host studies is that replication mode and intracellular constraints leave recognizable signatures in the accumulation of genomic strands, mutational spectra, and the fate of minority variants. Nested theoretical models that couple viral kinetics and immunity predict that infection outcomes (acute *vs* chronic), the strength and timing of selection, and the opportunities for variant establishment all emerge from the interaction between viral replication and host immune responses, rather than being fixed attributes of a pathogen (Alizon and van Baalen 2008). At the same time, spatial structure within tissues (*i.e*., limited dispersal distances, localized innate responses and heterogeneous cell states) can slowly spread, induce genetic surfing, and modulate the coupling between viral load and diversity, with important consequences for the interpretation of strand-specific dynamics and variant filtering (Gallagher et al. 2018). Recent syntheses underscore that capturing both transmission routes (cell-free and cell-to-cell) and spatial constraints is essential for realistic within-host models of viral spread (Dobrovolny 2025).

The host environment is a major component of this filtering process. Developmentally driven changes in cell identity, metabolism and immune competence create temporally shifting niches for replication and selection, thereby altering when and where diversity is generated and filtered. For example, plant and animal systems show that viral infections reprogram host primary metabolism and developmental patterns in ways that can sustain replication or shape defense, emphasizing that the intracellular environment is not static (Melero et al. 2023; Castiglioni et al. 2025; Melero et al. 2025). Similarly, the organization of tissue microbiota and local immune ecosystems across development can influence viral tropism, replication efficiency, and the strength of selective filters acting on viral populations (de Steenhuijsen Piters et al. 2015). Beyond developmental stage, host aging also plays a crucial role in pathogen evolution (Ben-Ami 2019). Thus, even when the focus is on viral replication dynamics, the intracellular environment encountered by the virus should be viewed as changing over the course of infection rather than static.

The Orsay virus (OrV) - *Caenorhabditis elegans* system offers a tractable model to study these processes. OrV is a non-enveloped, positive (+)-sense, single-stranded, bisegmented RNA virus that establishes chronic, low-virulence intestinal infections in *C. elegans*, allowing high-resolution tracking of replication and diversification over short timescales (Félix et al. 2011; Herek et al. 2026). RNA1 encodes the RNA-dependent RNA polymerase (RdRP), whereas RNA2 encodes the capsid protein (CP), the δ protein, and, through a ribosomal frameshift, a CP-δ fusion protein. CP-δ is necessary for cell attachment and entry (Fan et al. 2017), and δ is required for the nonlytic egress of new particles (Yuan et al. 2018). OrV, together with Santeuil (SANTV), Le Blanc (LEBV), and Mělník (MELV) form a clade of nematode-infecting noda-like viruses (Frézal et al. 2019).

Here, we exploit dense temporal, strand-specific RNA-seq data generated from synchronized OrV infections (Castiglioni, Olmo-Uceda, Villena-Giménez et al. 2024) to quantify (+)- and minus (−)-strand accumulation, characterize strand-resolved nucleotide diversity, identify strand-restricted mutations, and detect non-standard viral genomes (nsVGs). We hypothesized that strand-asymmetric replication generates variation in (−)-strand intermediates, only a subset of which is retained in (+)-strand genomes, thereby producing sequential filtering of both point mutations and structural variants. We tested this hypothesis progressively by examining RNA accumulation, global and local diversity, cross-strand SNV detection, functional constraint among shared variants, and recurrent nsVG formation.

## RESULTS

### Strand-asymmetric RNA accumulation reveals template-constrained replication

Previous work identified four phases of OrV accumulation during infection (Castiglioni, Olmo-Uceda, Villena-Giménez et al. 2024). Here, we quantified the (+)- and (−)-strand RNA species of both genomic segments across this time course (Fig. 1A).

**Figure 1.**
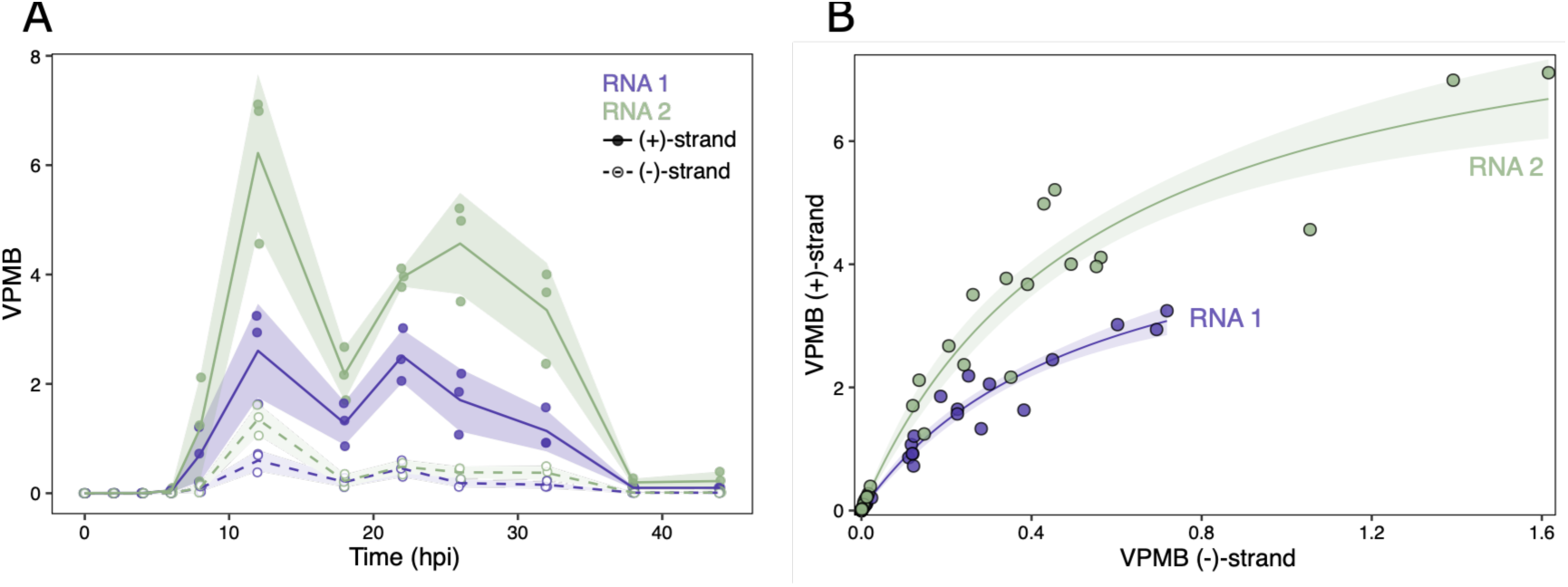
OrV accumulation. **(A)** Accumulation of viral RNA species over the course of infection. Solid lines and filled dots: positive strands; dashed lines and empty dots: negative strands. Lines are connecting the mean ±1 SD of the three biological replicates per timepoint. **(B)** Dependence of (+)-strand on (–)-strand abundance. The relationship was fitted to the saturating function (*i* = RNA1, RNA2): *RNA_i_*^(+)^= *a_i_RNA_i_*^(-)^/(*b_i_*+*RNA_i_*^(-)^), where *a_i_* is the asymptotic maximum abundance of (+)*RNA_i_*, and *b_i_* the abundance of (−)*RNA_i_* at which the fitted (+)RNA abundance reaches half of its asymptotic maximum. Purple: RNA1; green: RNA. VPMB, viral genomes per million of sequenced bases.

To test for global differences in accumulation between RNA segments, time post-inoculation, and RNA strand polarity, we fitted a generalized linear mixed model (GLMM) to the data in Fig. 1A, with sample as a random effect. RNA2 accumulated significantly more than RNA1 [mean effect = 0.35 ±0.06 (±1 SE), *P* < 0.001], and (+)RNAs exceeded (−)RNAs (2.31 ±0.06, *P* < 0.001). Time also had a significant positive effect (0.10 ±0.03, *P* = 0.002).

Next, we examined the proportions of the four viral molecular species throughout the course of infection (Fig. 1A). Comparisons between consecutive time points were used to assess progressive changes, whereas comparisons with the initial distribution (0 hpi) were used to evaluate overall divergence from the origin. All comparisons were significant (Table S1; *P* ≤ 0.016). The largest shift occurred between 22 and 26 hpi, when the abundance of all species declined except for (+)RNA2, which continued to increase (Fig. 1A; Table S1). The time point that differed most from the initial distribution was the final sampling point (Table S1). Together, these results indicate that differences among RNA species were reflected not only in their abundance but also in their relative contributions throughout infection (Fig. 1A, Fig. S1).

Because changes in relative proportions may arise from differences in accumulation dynamics, we quantified the similarity of temporal profiles using dynamic time warping (DTW). The two (−)RNA profiles were the most similar (distance = 0.09 ±0.01), whereas the (+)RNA profiles differed substantially (0.63 ±0.09). Within genomic segments, strand-to-strand DTW distances were greater for RNA2 (0.66 ±0.06) than for RNA1 (0.32 ±0.04). These differences can be summarized as follows: (*i*) (+)RNA2 exhibited a delayed second wave of accumulation, reaching its maximum abundance later than the other species and showing a correspondingly delayed decline; and (*ii*) when comparing the two accumulation peaks (12 and 22 hpi for (+)RNA1 and (−)RNA2, and 12 and 26 hpi for (+)RNA2), (+)RNA1 reached similar levels at both peaks, whereas the second peak was lower for (+)RNA2 and for the (−)-strands of both segments.

Collectively, these results indicate that strand- and segment-specific accumulation patterns became partially uncoupled during the second phase of infection, largely as a consequence of the delayed accumulation and decline of (+)RNA2.

(+)- and (−)-strand abundances were strongly correlated for both genomic segments (Fig. 1B; Spearman’s *r_S_* = 0.985 for RNA1 and *r_S_* = 0.980, for RNA2, 34 d.f., *P* < 0.001 in both cases). However, the relationship was nonlinear and was best described by a saturating model. Nonlinear least-squares fitting yielded *a_RNA1_* = 5.4 ±0.6, *a_RNA2_* = 9.0 ±0.8, *b_RNA1_* = 0.5 ±0.1, and *b_RNA2_* = 0.6 ±0.1 (all *P* < 0.001). For both genomic segments, the saturating model provided a substantially better fit than an exponential model, with lower AIC values, smaller residual variance, and Akaike weights strongly favoring saturation (Table S2). Together, these results show that OrV RNA accumulation is both strand- and segment-dependent and that (+)-strand abundance increases nonlinearly with (−)-strand abundance.

### Viral diversity is higher in (−)-strands and concentrated in specific genomic regions

Normalized Shannon entropy differed among molecular RNA species (Fig. 2A). For (+)RNAs, entropy increased and stabilized from 6 hpi onward, whereas (−)RNAs reached their maxima around 18 hpi. To assess effects of RNA segment, time post-inoculation, and strand on *S_n_*, we fitted a linear mixed model (LMM) with a random intercept for each sample. RNA2 had significantly lower entropy than RNA1 (effect = −0.002 ±0.001, *P* = 0.027), entropy increased with time [effect = (1.6 ±0.5)×10^−4^, *P* = 0.002], and (−)RNAs showed marginally higher entropy than (+)RNAs (effect = −0.002 ±0.001, *P* = 0.050).

**Figure 2.**
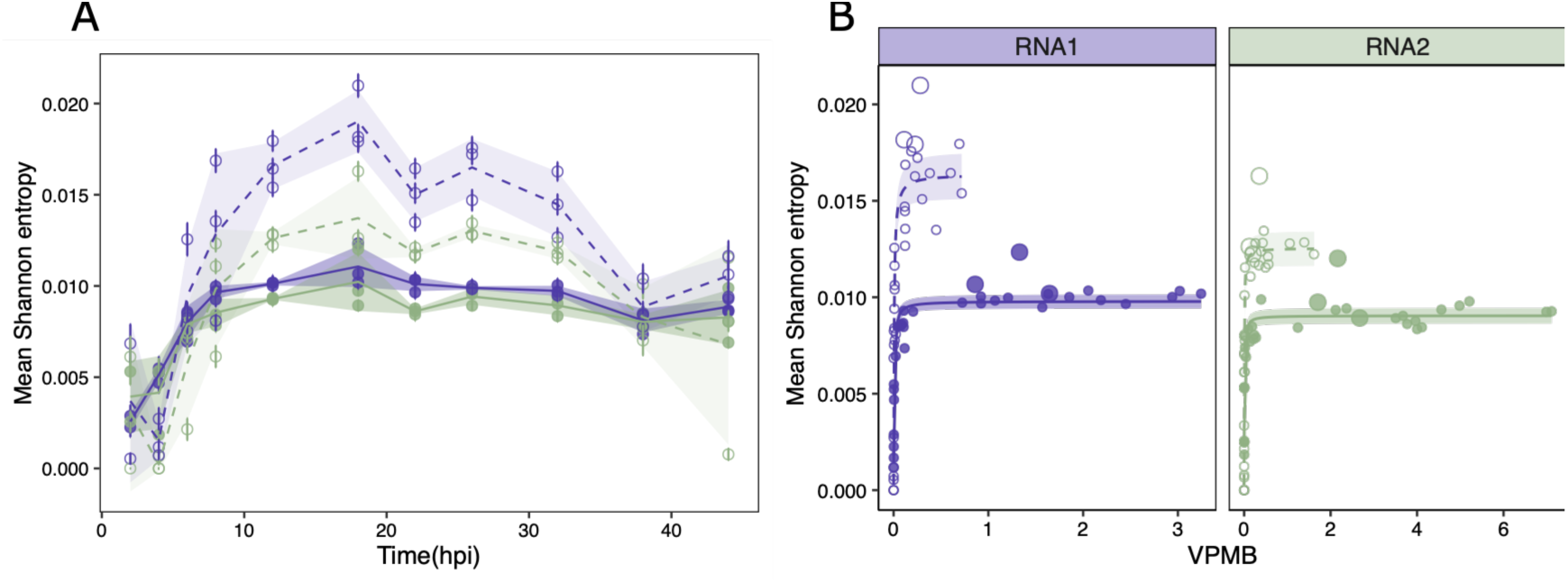
Evolution of OrV diversity during infection. **(A)** Dynamics of mean normalized Shannon entropy, *S̄_n_*. Dots show *S̄_n_* ±SEM for individual samples, lines connect the mean ±1 SD of the three replicates means per timepoint and viral specie. **(B)** Relationship between *S̄_n_* and viral accumulation (VPMB). Lines represent fitted nonlinear saturation models and shaded areas indicate 95% confidence intervals of the fitted curves (see footnote Table 1 for the model equation). In both panels, purple denotes RNA1 and green denotes RNA2; filled dots and solid lines indicate (+)RNAs while empty dots and dashed lines indicate (−)RNAs.

**Table 1.**
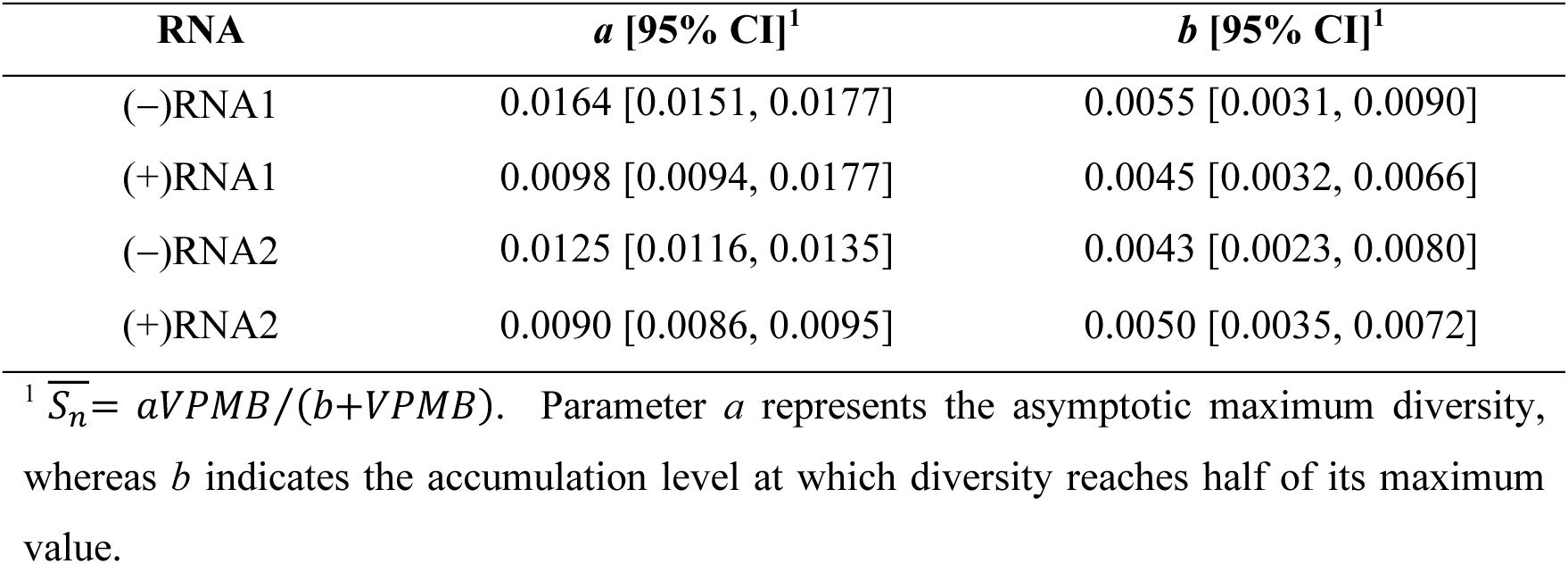
Estimated parameters of the saturation model relating mean nucleotide diversity (*S̄_n_*) to viral accumulation (VPMB), fitted separately for each RNA segment and strand polarity. Confidence intervals (CI) correspond to 95% intervals derived from the nonlinear least-squares model.

Notably, at 18 hpi especially (−)-strand species reached their maximum diversity while overall viral accumulation underwent a marked decline (Fig. 1A). This observation prompted an explicit analysis of the association between accumulation and diversity.

The relationship between viral accumulation and diversity was consistently described by a two-parameters saturating function across all viral species (Fig. 2B; Table 1). In every case, mean diversity increased rapidly at low VPMB values but approached a well-defined maximum asymptotic value *a* as accumulation increased, indicating the presence of an upper bound to sequence variability. The *a* differed significantly between strands and, to a lesser extent, between segments. Bootstrap comparisons confirmed that (−)RNAs reached significantly higher plateau values than (+)RNAs in both RNA segments (Table 1, Table S3). Additionally, (−)RNA1 exhibited significantly higher maximum diversity than (−)RNA2, whereas no differences were detected between (+)RNAs segments. Despite these differences in maximum diversity, no significant differences were observed in the VPMB value at which half of the maximum asymptotic diversity was achieved, *b*, across any comparison (Table S3), indicating that the accumulation range over which diversity approaches saturation is broadly conserved.

Thus, the accumulation range over which diversity approached saturation was similar among RNA species, whereas maximum diversity was consistently higher for (−)RNA. Position-resolved diversity was also strand-asymmetric. A pattern of diversity hotspots emerged at 6 hpi and persisted through the final time point (Fig. S1) and a LMM with position and sample as random effects showed that (−)RNA diversity tends to exceed that of (+)RNA (RNA1: effect = −0.0037 ±0.0002, *P* < 0.001; RNA2: effect = −0.0013 ±0.0002, *P* < 0.001).

We then asked which positions show the largest strand-dependent diversity differences. Because time had no overall significant effect in the LMM, we used the position-wise residual average to quantify strand bias at position *p* (Fig. 3). Positions with residual values above the 99^th^ percentile or below the 1^st^ percentile were considered strongly strand-biased and are highlighted in purple in Fig. 3A. Next, a 50-nt sliding window with 20-nt steps was applied to smooth the position-wise residual signal and identify genomic regions enriched in strand-biased diversity. The sliding-window results (Fig. 3B, middle) reveal that these enriched regions in the (−)RNA are concentrated in the final part of the RNA1 5’UTR, the end of RdRP, and the distal end of the first third of the CP and δ genes. In contrast, positions biased toward higher diversity in the (+)RNA strand were mainly restricted to the beginning of the 5’UTR in both genomic segments.

**Figure 3.**
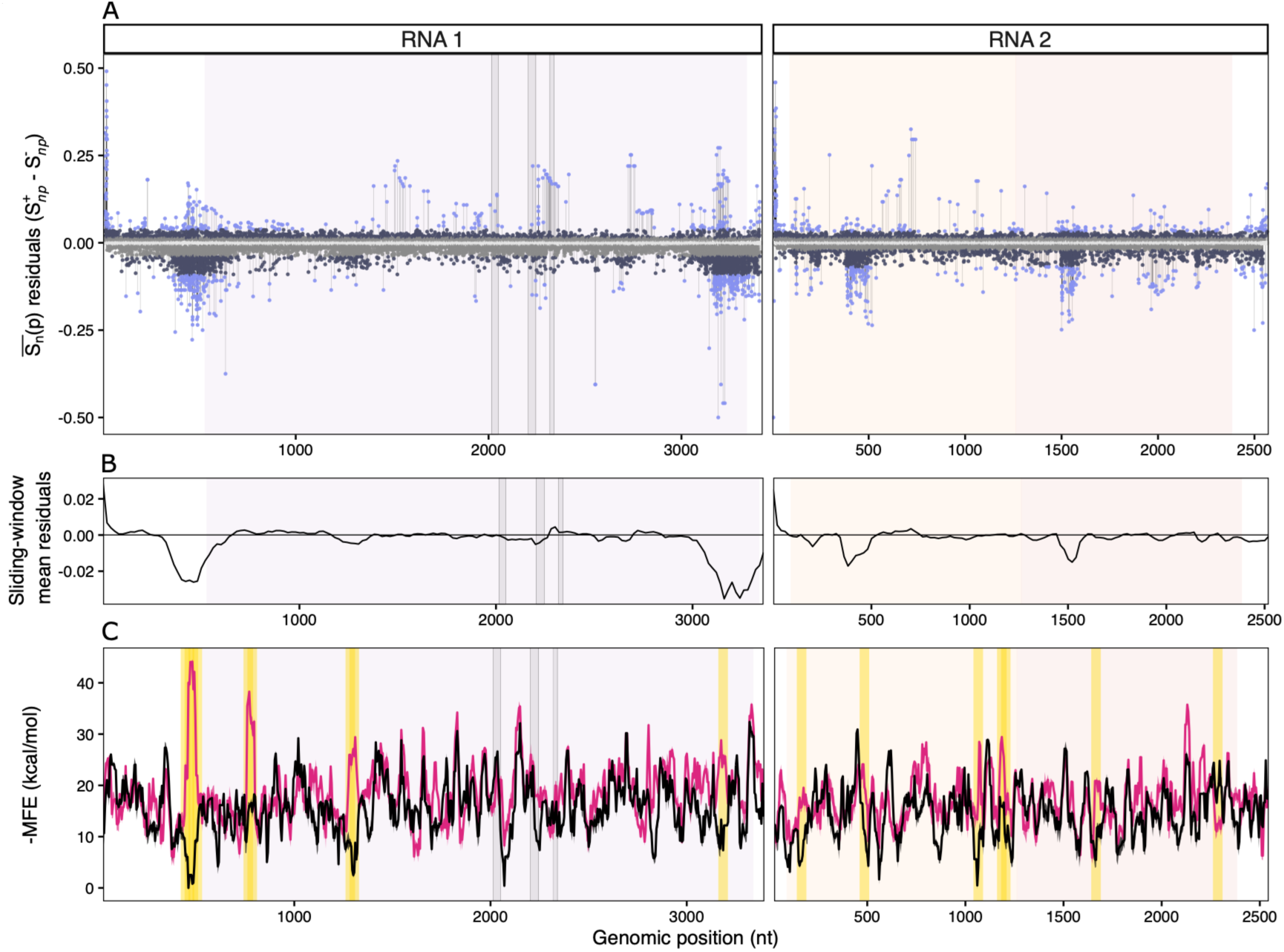
**(A)** Position-wise residual mean of nucleotide diversity differences between genomic strands, calculated as Δ*S̄_n_*(*p*)=*S̄_n_*^+^(*p*)-*S̄_n_*^-^ (*p*). Positive values indicate higher diversity in (+)RNAs, whereas negative values indicate higher diversity in (−)RNAs. Points were classified according to segment-specific empirical distributions of residual values: central (25^th^ - 75^th^ percentile, not plotted), moderate (outside interquartile range, in light gray), strong (outside 5^th^–95^th^ percentile, dark gray), and extreme (outside 1^st^ - 99^th^ percentile, purple). **(B)** Sliding-window average of strand-specific diversity residuals calculated using 50-nt windows and 20-nt steps. Negative values identify genomic regions where nucleotide diversity preferentially accumulates in the (−)-strand. **(C)** Predicted RNA structural potential for (+) and (−) strands along both genomic segments. Minimum free energy (MFE) was estimated with RNAfold using 50-nt sliding windows with 5-nt steps, allowing G-quadruplex formation, and plotted as −MFE values, where larger values indicate higher predicted structural stability. Gold-shaded regions indicate windows with strong strand-specific structural asymmetry, defined independently for each segment as windows whose absolute structural bias exceeded the segment-specific 95^th^ percentile. Per position *p* structural bias was calculated as Δ*MFE*(*p*)=*MFE*^(+)^(*p*)-*MFE*^(-)^(*p*), such that negative values indicate greater predicted secondary structure stability in the (−)-strand. In all the plots, coding regions are shown as shaded colored boxes (in order RdRP, CP and δ) and conserved RdRP palmprint motifs as grey regions. **(D)** Distribution of *S_n_* along the viral genome. Per site *S_n_* was calculated independently for each strand; only positions with > 10 reads were included. For clarity, sites with *S_n_* = 0 are uncolored. Dot and segment colors range from low entropy (yellow) to high entropy (purple). Background shading indicates coding regions. Gray rectangles mark the A, B, and C motifs of the RdRP palmprint domain.

We next asked whether strand-dependent diversity patterns could be associated with local differences in predicted RNA structural potential (Fig. 3C). RNAfold was run allowing G-quadruplex formation; therefore, MFE values should be interpreted as reflecting both canonical secondary structures and G-quadruplex-compatible conformations. This distinction was particularly relevant for the most asymmetric region in RNA1, where the corresponding (+)-strand window was C-rich and weakly structured, whereas the reverse-complement (−)-strand window was G-rich and predicted to form highly stable G-quadruplex-compatible structures, suggesting that extreme structural asymmetry may contribute to local strand-dependent diversity patterns. To quantify this relationship genome-wide, we compared strand-dependent diversity residuals with structural asymmetry, calculated as the difference in predicted MFE across matched sliding windows. Across all windows, we observed a weak but significant positive association between diversity residuals and structural asymmetry (Spearman’s *r_S_* = 0.17, 294 d.f., *P* = 0.003), indicating that regions with greater predicted structural differences between strands tended to display stronger strand-specific diversity shifts. This relationship was more pronounced in RNA1 (*r_S_* = 0.38, 167 d.f, *P* < 0.001), whereas RNA2 exhibited a weaker association in the opposite direction (*r_S_* = −0.19, 125 d.f, *P* = 0.034), suggesting that the coupling between predicted structural potential and diversity is not uniform across the genome. Permutation analysis confirmed that the observed global correlation was unlikely to arise by chance (*P* = 0.003), supporting a non-random association between structural and diversity asymmetries.

To further assess whether this relationship was driven by extreme regions, we defined structural hotspots as windows exceeding the 95^th^ percentile of absolute structural asymmetry within each segment. These hotspots, highlighted in yellow in Fig. 3C, showed significantly higher structural bias compared to non-hotspot regions (Wilcoxon test, *P* = 0.039). However, diversity hotspots did not show a corresponding enrichment in structural asymmetry (*P* = 0.136), and directional hotspot enrichment tests were not significant. Thus, predicted RNA structural potential, including G-quadruplex-compatible regions, may contribute to strand-dependent diversity patterns, but is not sufficient to explain all localized diversity peaks.

Overall, structural and diversity asymmetries were only modestly associated, and structural hotspots did not consistently overlap diversity hotspots. Predicted RNA structure therefore explained only a limited fraction of the spatial organization of strand-biased diversity.

### Strand-resolved SNVs reveal incomplete retention of viral variation

#### Negative strands contain greater SNV richness and abundance

Next, we called SNVs for all four viral molecular species. A preliminary summary of the number of SNVs identified in each segment is shown in Fig. S2A. Richness was computed as the number of SNVs (Fuhrman et al. 2021) above the AF threshold in each sample, irrespective of strand. We performed a Poisson GLMM with time and RNA segment as fixed effects, a log-offset for segment length (in order to control the differences in length of both segments), and sample as random intercept. No effect of time post-inoculation (effect = 0.01 ±0.01, *P* = 0.448) or RNA segment (RNA2 *vs* RNA1: effect = −0.002 ±0.074, *P* = 0.983) was detected. SNV abundance (Fig. S2B), defined as the cumulative AF of variants with AF > 0.01, was analyzed using an analogous Gamma GLMM with a log link. Abundance increased significantly over time (effect = 0.025 ±0.007, *P* < 0.001), whereas no difference was detected between RNA segments (RNA2 *vs* RNA1: effect = −0.1 ±0.1, *P* = 0.619).

Strand-resolved analyses were more informative (Fig. 4). Incorporating strand into the models did not reveal any effect of time or between segments on richness (hpi: effect = 0.01 ±0.01, *P* = 0.592; RNA2 *vs* RNA1: effect = 0.09 ±0.06, *P* = 0.139). However, the (+)-strand exhibited significantly lower SNV richness than the (−)-strand (effect = −0.15 ±0.06, *P* = 0.016), corresponding to an estimated 14% reduction. Similarly, SNV abundance (Fig. 4B) remained unrelated to RNA segment (RNA2 *vs* RNA1: effect = −0.033 ±0.098, *P* = 0.736) but increased significantly with time (effect = 0.015 ±0.006, *P* = 0.008), and was 22% lower on the (+)- than on the (−)-strand (effect = −0.2 ±0.1, *P* = 0.009).

**Figure 4.**
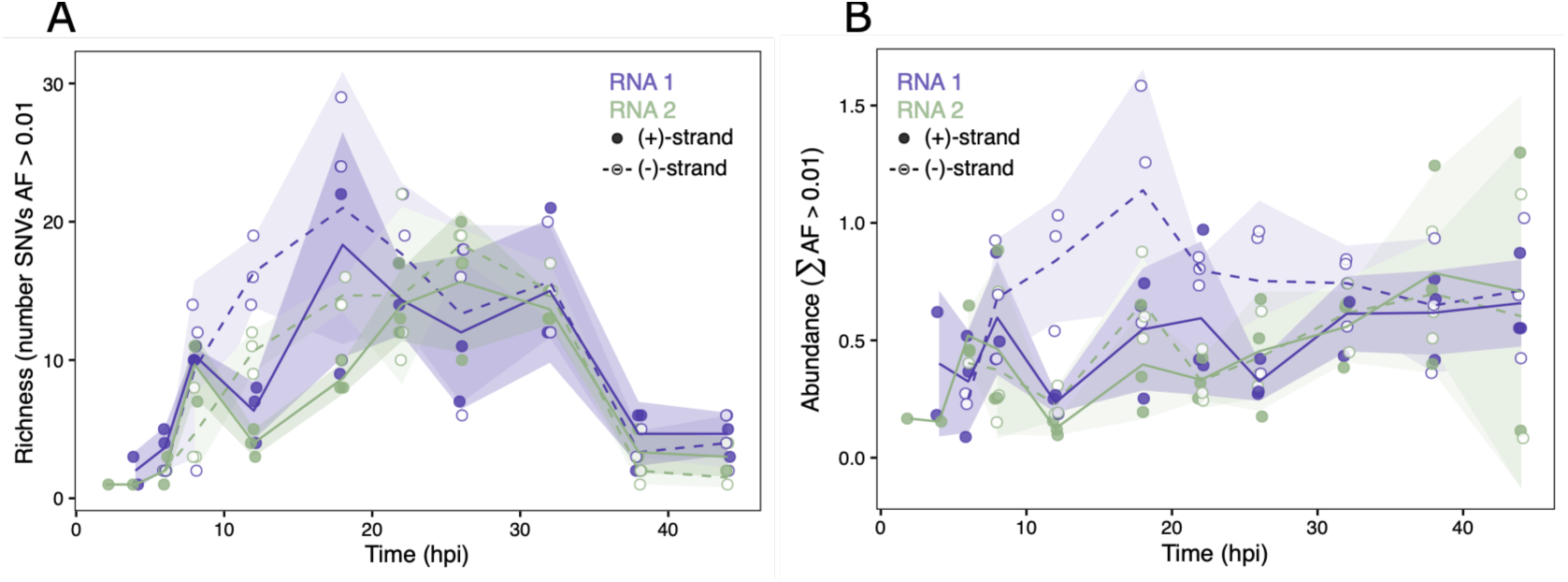
SNV richness and accumulated allele frequency (AF). **(A)** SNV richness dynamics. Richness as the number of SNVs with AF > 0.01 of each viral specie per sample. **(B)** SNV abundance dynamics. Abundance is the sum of AF across all SNVs with AF > 0.01. In both panels, lines show the mean profile for positive (solid) and negative (dashed) strands (shadow indicates ±1 SD; individual points for each sample values are solid for (+) and empty for (−) strand. RNA1 in purple and RNA2 in green.

These results suggest two, not mutually exclusive, possibilities. (*i*) Diversity peak and second accumulation wave. The diversity peak at 18 hpi could reflect immune escape that enables a second accumulation wave. Alternatively, because signs of egress were detected as early as 14 hpi (Castiglioni, Olmo-Uceda, Villena-Giménez et al. 2024), the peak may represent variants with slower replication cycles: as fast-replicating variants complete replication, slower variants (potentially carrying more mutations) could finish their cycles and transiently dominate. (*ii*) Within-cell constraint on (+)-strands. The presence of SNVs on (−)RNAs without corresponding signals on (+)RNAs could indicate a within-cell constraint or filtering process that limits the propagation of certain mutations to positive strands. These hypotheses are explored in the following section.

#### Only a subset of (−)-strand SNVs is detected in (+)-strands

SNVs were classified within each sample as (+)RNA-specific, (−)RNA-specific, or shared between strands (Fig. S3A-B). Only variants with AF > 0.01 were considered. We then quantified conditional strand overlap, defined as the proportion of SNVs detected on (−)RNA that were also detected on (+)RNA (Fig. S3C). A substantial fraction of variants detected on (−)RNA was not detected on (+)RNA during early infection, whereas cross-strand detection increased later. Temporal variation in propagation probabilities was evaluated separately for each genomic segment using quasibinomial GLMs, and pairwise comparisons among time points on the log-odds scale using estimated marginal means. For RNA1, the estimated conditional probability exhibited its minimum at 12 hpi [0.245, 95% CI = 0.139 - 0.395], indicating that fewer than one quarter of variants detected on (−)RNA were also observed on (+)RNA. Propagation probability increased thereafter, reaching 0.540 [95% CI = 0.408 - 0.666] at 18 hpi and 0.642 [95% CI = 0.495 - 0.766] at 22 hpi. The log-odds of overlap at 12 hpi was significantly lower than at 18 hpi (odds ratio (OR) = 0.277, adjusted *P* = 0.049), 22 hpi (OR = 0.181, adjusted *P* = 0.010), and 32 hpi (OR = 0.240, adjusted *P* = 0.049) (Fig. S3C). For RNA2, although statistical support was weaker and none of the pairwise contrast remained significant after multiple-testing correction, the same temporal pattern was observed, with the lowest estimated transmission probabilities consistently occurring around 12 hpi. This pattern is consistent with apparent within-cell filtering, but does not distinguish among differential retention, intracellular bottlenecks, strand-specific detectability, degradation, and stochastic sampling.

#### Strand-specific SNVs are spatially clustered

We then examined the genomic distribution and recurrence across samples to assess whether strand-specific SNVs were randomly distributed along each genomic segment. Kernel density estimates (KDE) weighted by the number of samples carrying each SNV were compared against a null distribution generated from 5,000 random permutations of SNV positions while preserving the number of variants and their recurrence weights. Genomic regions where the observed density exceeded the 97.5^th^ percentile of the null distribution were considered enriched for strand-specific SNVs (Fig. 5). The results suggest that the C-terminal domain of RdRP is especially enriched for such events. In addition, the CP variants 231U>C (I48T), 237U>A (L50Q) and 243G>A (G52D) were detected in five, ten and six samples respectively, but only on the (−)RNA (Fig. 5). SNVs at positions 2199, 2205, 2206 (P316T), and 2212 (N318D) within the δ protein were identified in at least 11 samples, again only on the (−)-strand.

**Figure 5.**
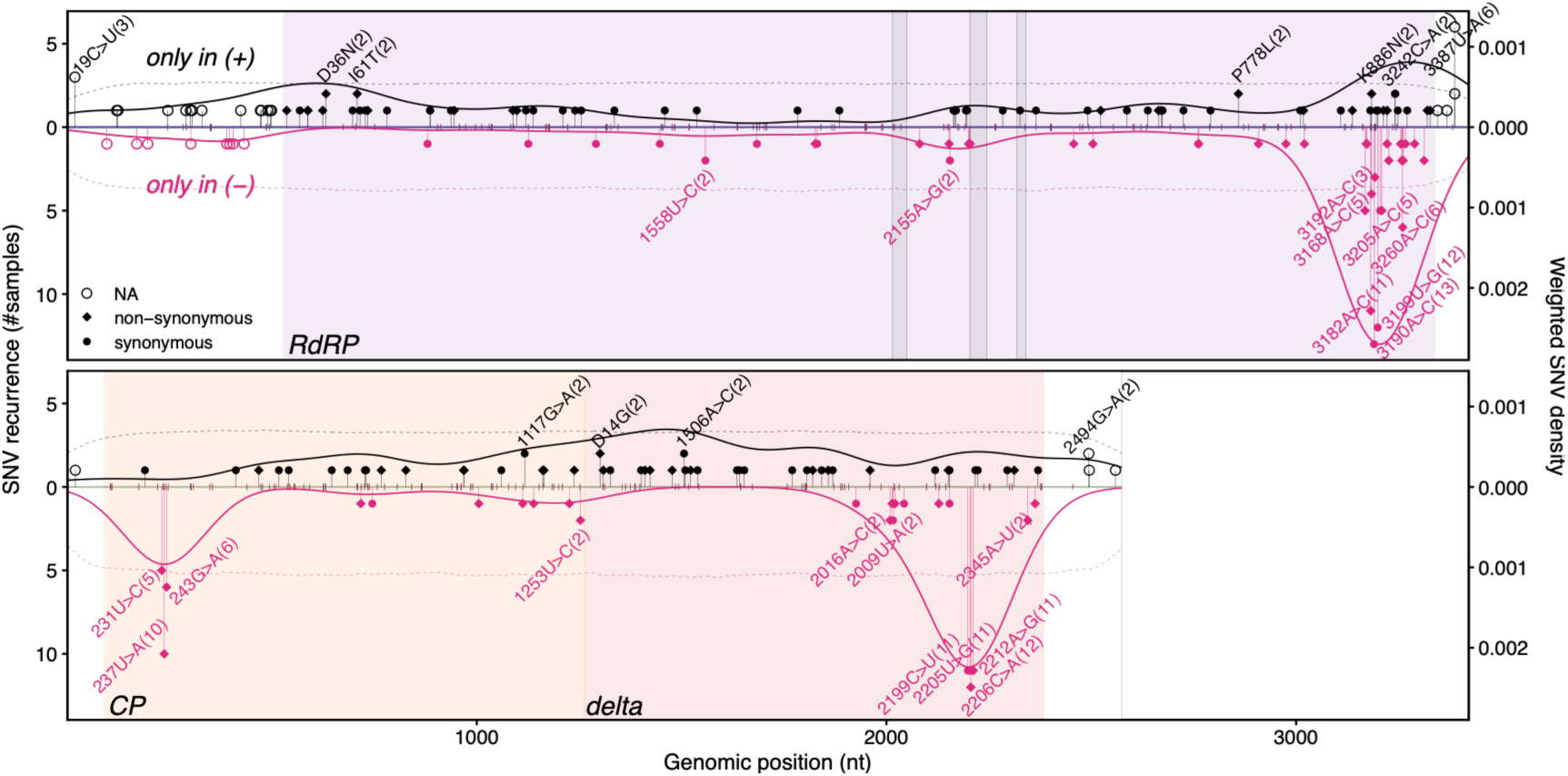
Distribution of strand-specific SNVs along OrV genome. SNVs detected exclusively on one strand are plotted at their genomic positions; segment height indicates the number of samples in which the SNV was independently observed. (+)RNA-specific SNVs are shown in black and (−)RNA-specific SNVs in pink. Burgundy short segments mark the position of the SNVs found in both strands without representing recurrence or density information. Background colors denote coding regions, consistent with other figures. Labeled SNVs are those detected in more than one sample (some labels are omitted in especially dense regions). Labels show the nucleotide change for noncoding or synonymous variants and the amino acid change for nonsynonymous variants [only in (+)RNA-specific SNVs]. The number of samples in which an SNV appears with AF > 0.01 is given in parentheses. Point shapes indicate the functional effect of the change, as defined in the legend. Solid curves above (black) and below (pink) the zero line represent the weighted kernel density of (+)RNA-specific and (−)RNA-specific SNVs, respectively. Dashed curves indicate the corresponding upper 95% CI obtained from 5,000 random permutations of SNV positions while preserving SNV recurrence. Regions where the observed density exceeds this threshold indicate significant local enrichment of strand-specific SNVs.

The substitution spectrum found in these SNVs does not indicate a consistent global bias toward transitions over transversions, and the relative contribution of both classes varies across RNA segments and strands (Fig. S4A). We further examined whether strand-specific SNVs could reflect known RNA editing processes. Although individual substitutions compatible with ADAR- and APOBEC-associated signatures are present in the (+)-associated SNVs, their distribution is not coherent across strands or genomic regions and does not show enrichment in the regions of SNV accumulation (Fig. S4B).

The recurrent localization of strand-specific SNVs in discrete genomic regions was inconsistent with a uniform genomic distribution, although the present data cannot resolve the underlying mechanism.

#### Shared SNVs are AF concordant, recurrent and functionally constrained

We found that the same-sample cross-strand SNVs showed a strong correlation of their AFs (*r* = 0.927, 302 d.f., *P* < 0.001; Fig. 6A). A linear model including mutation effect revealed that non-synonymous variants exhibit reduced strand concordance compared to synonymous variants (slope = 0.750 *vs* 0.955; Δ = −0.21 ±0.04, *P* < 0.001). Extending the model to include proteins showed no overall effect of CDS on AF concordance, although a modest region-specific deviation in slope was observed in RdRP (*P* < 0.001 for interaction), indicating a slight reduction in strand concordance in this segment (Fig. 6A). Additionally, the proportion of synonymous *vs* nonsynonymous variants varied significantly across proteins (LRT: χ² = 10.51, 2 d.f., *P* = 0.005) driven mainly by RdRP, which showed an enrichment of synonymous variants relative to CP (pairwise contrast, *P* = 0.007), whereas differences with δ were not (*P* = 0.062). A Fisher’s exact test revealed a significant difference in the frequency of nonsynonymous variants between the genomic segments: RNA1 had a significantly lower proportion of nonsynonymous variants than RNA2 (Fig. 6A; OR = 0.43, *P* = 0.003).

**Figure 6.**
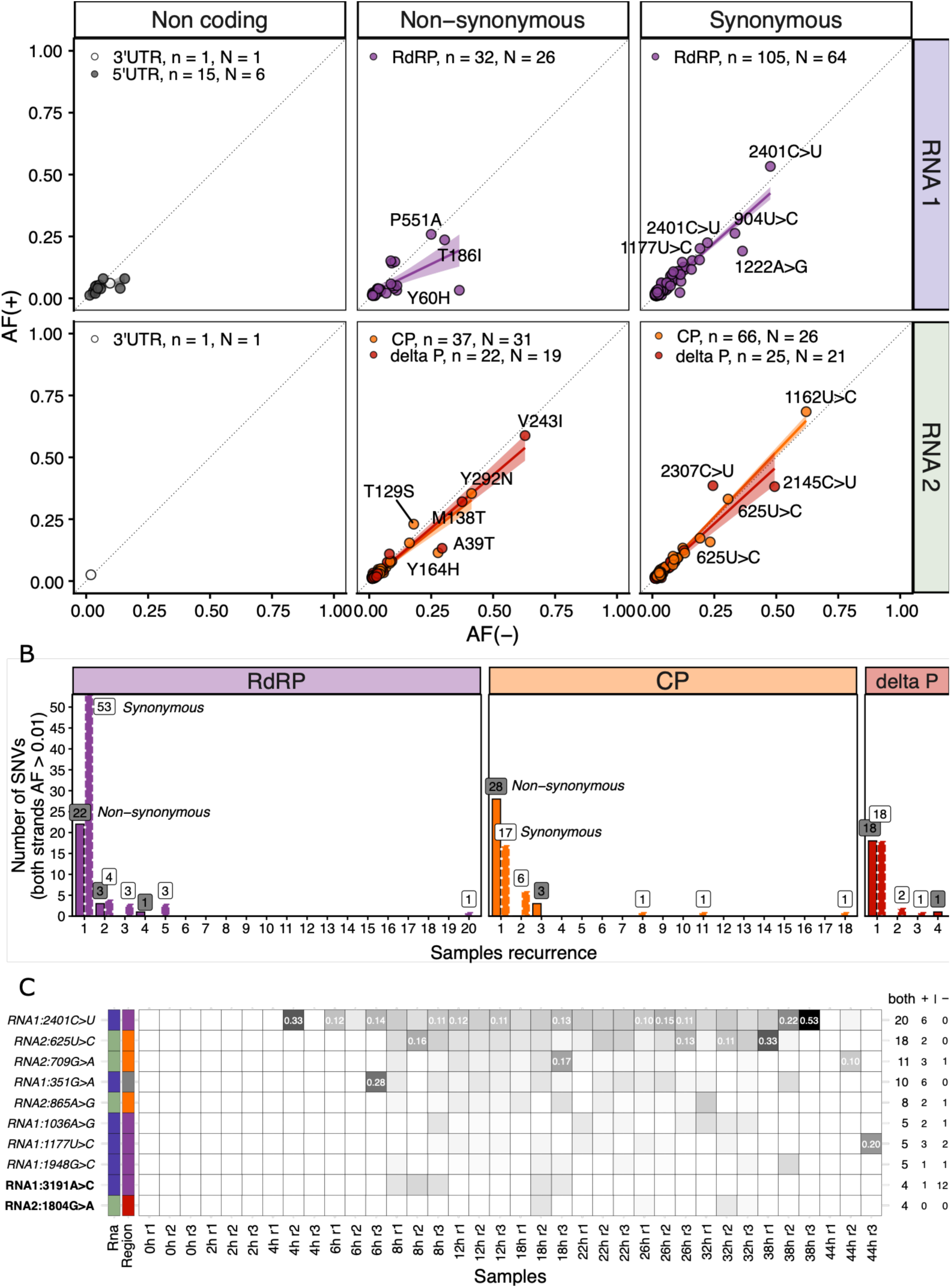
SNVs detected on both strands (with AF > 0.01). **(A)** Correlation of allele frequencies between (+)- and (−)-strands. The total number of SNVs (*n*) and the number of unique SNVs (*N*) per region and amino-acid effect are indicated in each panel. **(B)** Distribution of the number of samples in which each the SNVs was were detected in both strands recurrently. Colors denote proteins or UTRs as in panel A. Bars with solid border and label with gray background for nonsynonymous changes; bars with discontinuous border and white background label for synonymous changes. **(C)** Heatmap of SNVs found on both strands in at least four samples with AF > 0.01 in both strands. The intensity represents the AF in the (+)-strand from white (below the AF threshold) to black (AF = 0.53). SNVs with an AF(+) > 0.1 are indicated. The number of samples were the SNVs has been detected on both strands, only in the (+)- or in the (−)-strand above the AF threshold are indicated in the right side of the heatmap. On the left side the RNA segment and the genomic region affected are coded with colors as in (A) and (B). If the nucleotide variation results in a nonsynonymous amino acid substitution, the SNV is shown in bold; otherwise, it is shown in italics.

We then asked whether these SNVs were recurrent across samples. Synonymous variants showed a higher probability of recurring in at least two independent samples than nonsynonymous variants (LRT, OR = 2.43, *P* = 0.043), whereas no effect of viral protein was detected (Fisher’s test, *P* = 0.368). This difference was no longer significant when only variants recurring in three or more samples were considered (*P* = 0.395; Fig. 6B). Overall, 21.1% of SNVs present on both strands appeared in more than one sample. When grouped by protein and by amino acid effect, the percentages of recurrent SNVs (nonsynonymous *vs* synonymous) were: RdRP (13.3% *vs* 22.9%), CP (10.3% *vs* 37.0%) and δ (4.76% *vs* 16.7%) (Fig. 6B).

Finally, Fig. 6C shows the allele frequencies of all SNVs detected on both strands (AF > 0.01) at least four samples. Only two of these were nonsynonymous (bold in Fig. 6C): RNA1 mutation 3191A>C (K889Q) and RNA2 mutation δ 1804G>A (V182I). Notably, K889Q was the only mutation observed either on both strands or exclusively on the (−)RNA, where it appeared in 12 independent samples.

Although our sampling design does not permit lineage-level tracking, we found that recurrent SNVs were generally detected at AF higher than the sample-specific background mutational spectrum. Across independent samples between 8 and 32 hpi, recurrent variants consistently showed higher median AF than non-recurrent SNVs from the same libraries, with this difference remaining significant after multiple-testing correction in most samples (Table S4).

We next tested whether shared coding variation showed heterogeneous functional constraint across the genome.

The distribution of *d_N_* – *d_S_* differed significantly among the three viral proteins (Kruskal-Wallis test, χ² = 24.321, 2 d.f., *P* < 0.001; Fig. 7A). Pairwise comparisons (Wilcoxon rank-sum tests with Benjamini-Hochberg (BH) correction) showed that δ exhibited significantly higher *d_N_* – *d_S_* values than both CP (*P* < 0.001) and RdRP (*P* < 0.001), whereas CP and RdRP did not differ significantly between them (*P* = 0.730).

**Figure 7.**
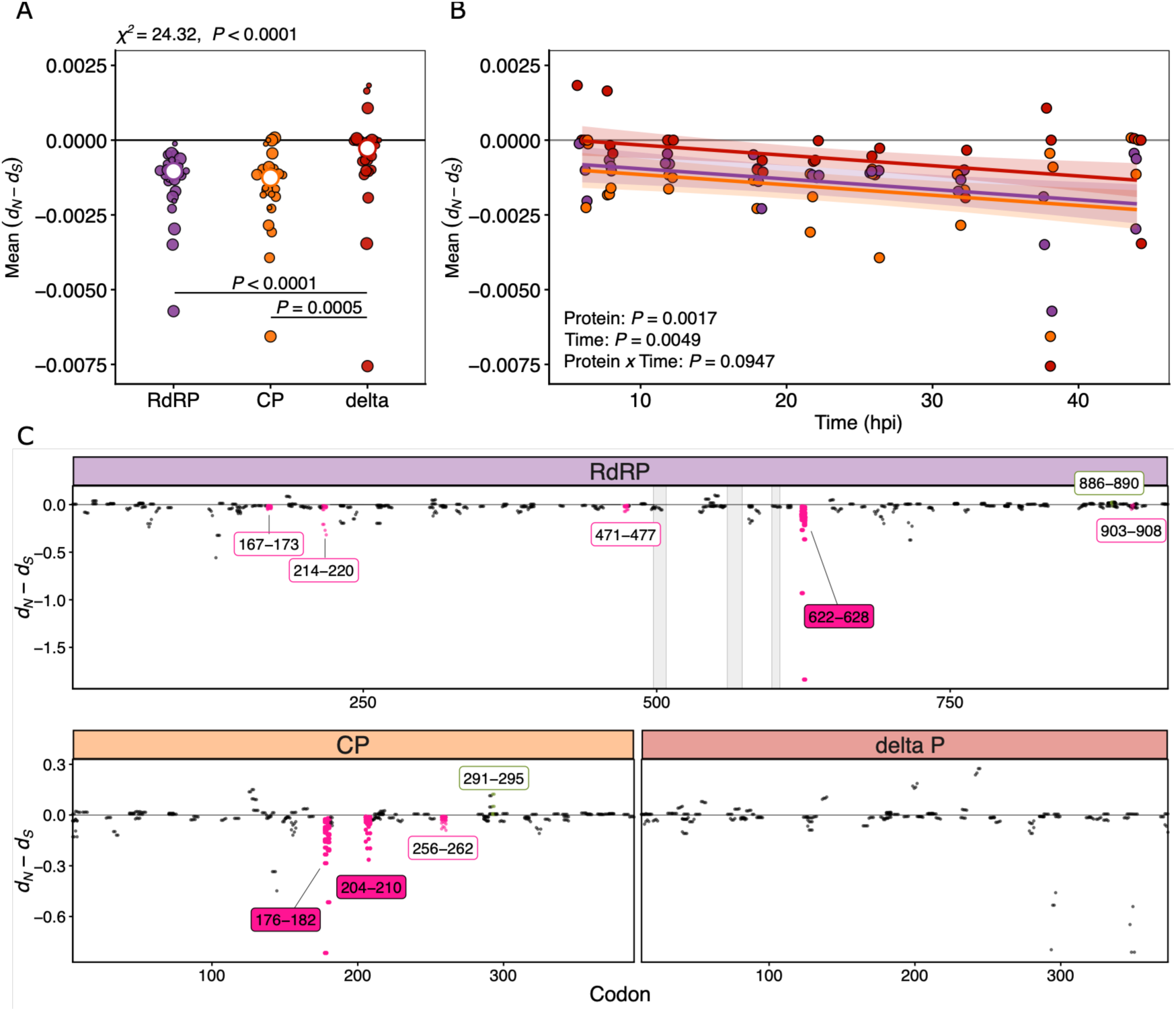
Global and local patterns of selection across OrV proteins. **(A)** Distribution of *d_N_* – *d_S_* values among viral proteins. Points represent the mean *d_N_* – *d_S_* of each biological sample (point size indicates hpi), although the analysis was performed with all the sample measures. Overall differences among proteins were assessed using a Kruskal-Wallis test (χ^2^ = 24.32, 2 d.f., *P* < 0.001) followed by BH-adjusted pairwise Wilcoxon rank-sum tests. *P* value is indicated when significant changes. **(B)** Temporal dynamics of sample mean *d_N_* – *d_S_*. Points correspond to biological samples, lines indicate protein-specific fit, with its 95% CI. Differences were evaluated using a linear mixed-effects model with protein and time as fixed effects and sample as a random effect. Adding the interaction (protein × time) did not improve the model and the interaction was not significant (*P* = 0.095). **(C)** Sliding-window analysis of *d_N_* – *d_S_* along each viral protein. Each point represents a 4-codon window. Windows significantly different from zero were identified using one-sided Wilcoxon signed-rank tests with BH correction applied separately for positive (green) and negative selection (magenta). Filled labels indicate significant windows after correction (adjusted *P* < 0.05) while non-filled labels indicate the windows significant before the correction (*P* < 0.05).

To determine whether selective pressure changed during infection, we then modelled window-specific *d_N_* – *d_S_* values using a linear mixed-effects model including protein and time as fixed effects and sample identity as a random intercept (Fig. 7B). Both protein (*P* = 0.003) and time post inoculation (*P* = 0.008) significantly affected *d_N_* – *d_S_*, whereas the protein × time interaction was not significant (*P* = 0.095). Estimated marginal means confirmed the observed overall differences among proteins. Both RdRP and CP showed significantly smaller *d_N_* – *d_S_* values than δ (adjusted *P* < 0.001 and adjusted *P* = 0.003, respectively), while CP and RdRP remained statistically indistinguishable (adjusted *P* = 0.440). In addition, we found that *d_N_* – *d_S_* tend to decreased as infection progressed [effect = (−3 ±1)×10⁻^5^ h⁻^1^, *P* = 0.005; Fig. 7B].

Finally, each sliding window was tested against the null hypothesis of *d_N_* – *d_S_* = 0 using one-sided Wilcoxon signed-rank tests. Three regions showed significantly negative *d_N_* – *d_S_* values (adjusted *P* < 0.05), whereas no window exhibited significantly positive cases (Fig. 7C). Mapping these regions onto the protein structures revealed that the negatively selected region in the RdRP (codon window 622-628) lies immediately adjacent to the conserved palm domain motifs and borders one of the major solvent-accessible channels of the polymerase structure (Fig. S5A-B). The two CP clusters (codon windows 176-182 and 204-210) are localized into the bottom-lateral face of the trimer capsid (Fig. S5C-F).

Notably, these three regions encompassed the recurrent synonymous SNVs identified in the previous result: RNA1 2401C>U (conserves A625), RNA2 625U>C (conserves the CP C179) and RNA2 709G>A (conserves the CP L207) (Fig. 6C, Fig. S5E) and two more SNVs that appear in three and two independent samples (RNA2 631U>C the CP G181 and RNA2 718C>U the CP A210) with an AF > 0.01 and in more below the selected threshold. Because the window-specific tests integrate *d_N_* – *d_S_* values across all samples, recurrent synonymous variants contribute consistently to the statistical support of these regions. The fact that the same variants were repeatedly detected in independently sampled infections, together with the non-random structural context of the corresponding regions, adjacent to conserved palm-domain motifs in RdRP and on the inner face of the CP, provides additional biological context for the localized patterns identified by the analysis.

### NsVG classes differ in their accumulation and genomic distribution

Because polymerase activity can generate both nucleotide substitutions and structural variants, we next examined whether OrV replication also produces recurrent classes of nsVGs. After clustering nearby junctions and requiring more than five supporting reads, 487 representative clusters were retained for analysis.

NsVG richness dynamics clearly reassemble the viral accumulation dynamics (Fig. 8A). To identify the factors associated with nsVG richness, we fitted a negative-binomial mixed-effects model including viral accumulation (VPMB), RNA segment and nsVG category as fixed effects, and sample identity as a random effect. Richness increased strongly with viral accumulation (effect = 2.7 ±0.2, *P* < 0.001). Inclusion of hpi did not significantly improve model fit after accounting for VPMB (LRT, *P* = 0.379), indicating that viral accumulation rather than infection time per se was the major determinant of richness. A significant interaction between RNA segment and nsVG category (LRT, *P* < 0.001) revealed segment-specific patterns of nsVG generation. In both RNA segments, deletions were the richest nsVG category, followed by insertions, whereas 3’ copy-backs (cb) and 5’cb nsVGs displayed substantially lower richness (all pairwise comparisons involving deletions, Tukey-adjusted *P* < 0.001). However, the magnitude of these differences depended on the RNA segment. While deletion richness differed only modestly between RNA1 and RNA2, insertion, 5’cb and 3’cb nsVGs exhibited markedly lower richness in RNA2, suggesting that the generation of non-deletion nsVGs may be constrained in a segment-dependent manner (Fig. 8A-B).

**Figure 8.**
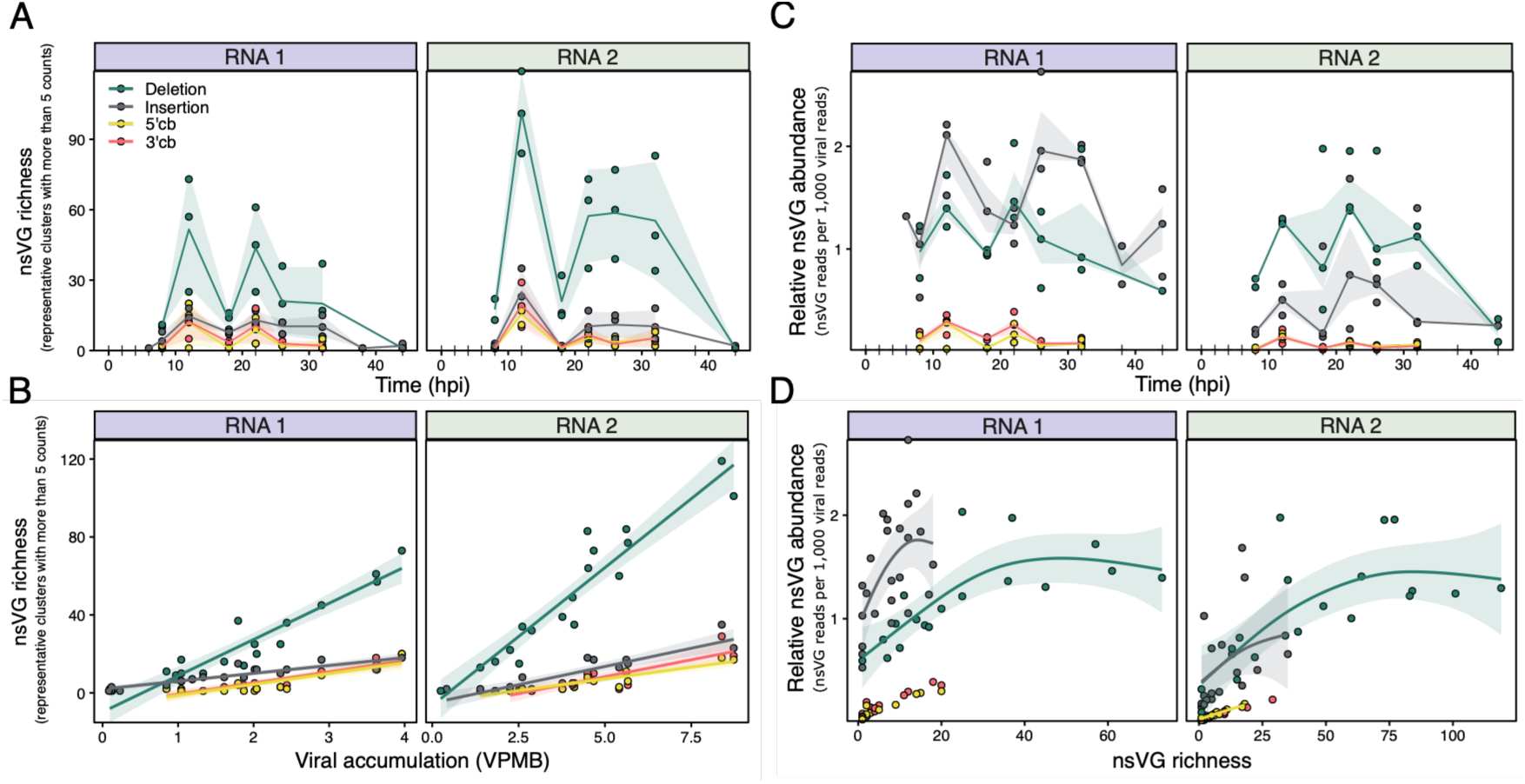
Dynamics and determinants of nsVG diversity during OrV infection. **(A)** Temporal dynamics of nsVG richness, defined as the number of representative nsVG clusters supported by > 5 reads, grouped by nsVG category and RNA segment. Points represent individual samples, lines correspond to the mean value at each time point, and shaded areas indicate ±1 SD. **(B)** Relationship between nsVG richness and viral accumulation (VPMB). Lines show linear model fits and shaded areas indicate SD. **(C)** Temporal dynamics of relative nsVG abundance, expressed as nsVG-supporting reads per 1,000 viral reads of the corresponding RNA segment. Points represent individual samples, lines correspond to mean values, and shaded areas indicate 95% CI. **(D)** Relationship between relative nsVG abundance and nsVG richness. Generalized additive models (GAMs) were fitted independently for each nsVG category. Shaded areas represent 95% CI. Colors indicate nsVG categories throughout the figure as in (A).

We next evaluated the relative abundance of nsVGs, expressed as the number of nsVG-supporting reads per 1,000 viral reads (Fig. 8C). Similar to richness, relative abundance was significantly associated with viral accumulation (*P* < 0.001), whereas hpi did not explain additional variation once VPMB was included in the model (LRT, *P* = 0.350). Relative abundance also differed significantly between RNA segments and nsVG categories (both *P* < 0.001), with a strong interaction between both factors (LRT, *P* < 0.001). Abundance patterns did not fully mirror richness patterns. In RNA1, insertion and deletion nsVGs reached the highest relative abundances (estimated marginal means = 1.04 and 0.85, respectively) and both accumulated significantly more than 3’cb and 5’cb nsVGs (Tukey-adjusted *P* < 0.001). In contrast, deletion nsVGs were the most abundant category in RNA2 (estimated marginal mean = 0.70), significantly exceeding insertion, 3’cb and 5’cb nsVGs (all Tukey-adjusted *P* < 0.001; Fig. 8C). Thus, although deletions consistently displayed the greatest richness, insertions contributed disproportionately to nsVG abundance in RNA1.

To further investigate the relationship between nsVG diversity and accumulation (Fig. 8D), we modelled relative abundance as a function of richness using generalized additive models. Allowing nsVG type-specific non-linear richness-abundance relationships significantly improved model fit compared with both a linear model (analysis of deviance, *P* < 0.001) and a model assuming a common richness-abundance relationship across nsVG categories (*P* < 0.001). Deletion and insertion nsVGs exhibited significant non-linear relationships, characterized by a rapid increase in abundance with increasing richness followed by progressively smaller gains at higher richness values. In contrast, both cb categories displayed approximately linear richness-abundance relationships, indicating a proportional increase in abundance as richness increased. Together, these results indicate that different nsVG classes are not only generated at different rates, but also differ in the efficiency with which increasing diversity translates into population-level accumulation during infection.

Consistent with these observations, analyses of overall nsVG community structure revealed that both effective Shannon diversity and community composition were primarily associated with viral accumulation rather than infection time (Fig. S6). Whereas effective Shannon diversity increased strongly with VPMB, community evenness showed a slight decline, indicating increasing dominance of a subset of nsVG clusters as viral accumulation increased (Supplementary Text 1).

Mapping representative clusters onto the viral genome revealed clear hotspots of nsVG formation for all four nsVG categories (Fig. 9). Deletions displayed the broadest genomic distribution and accumulated predominantly near the diagonal of the start-end plot, indicating that most deletion events involved the removal of relatively short genomic regions. However, several recurrent long-range deletion hotspots were also observed off diagonal. In RNA1, recurrent deletion clusters connected distant regions of the RdRP coding sequence, whereas RNA2 contained a prominent hotspot joining the 5’ region of the segment with the δ region. These long-range deletion clusters were among the most abundant deletion-derived nsVGs detected in the dataset.

**Figure 9.**
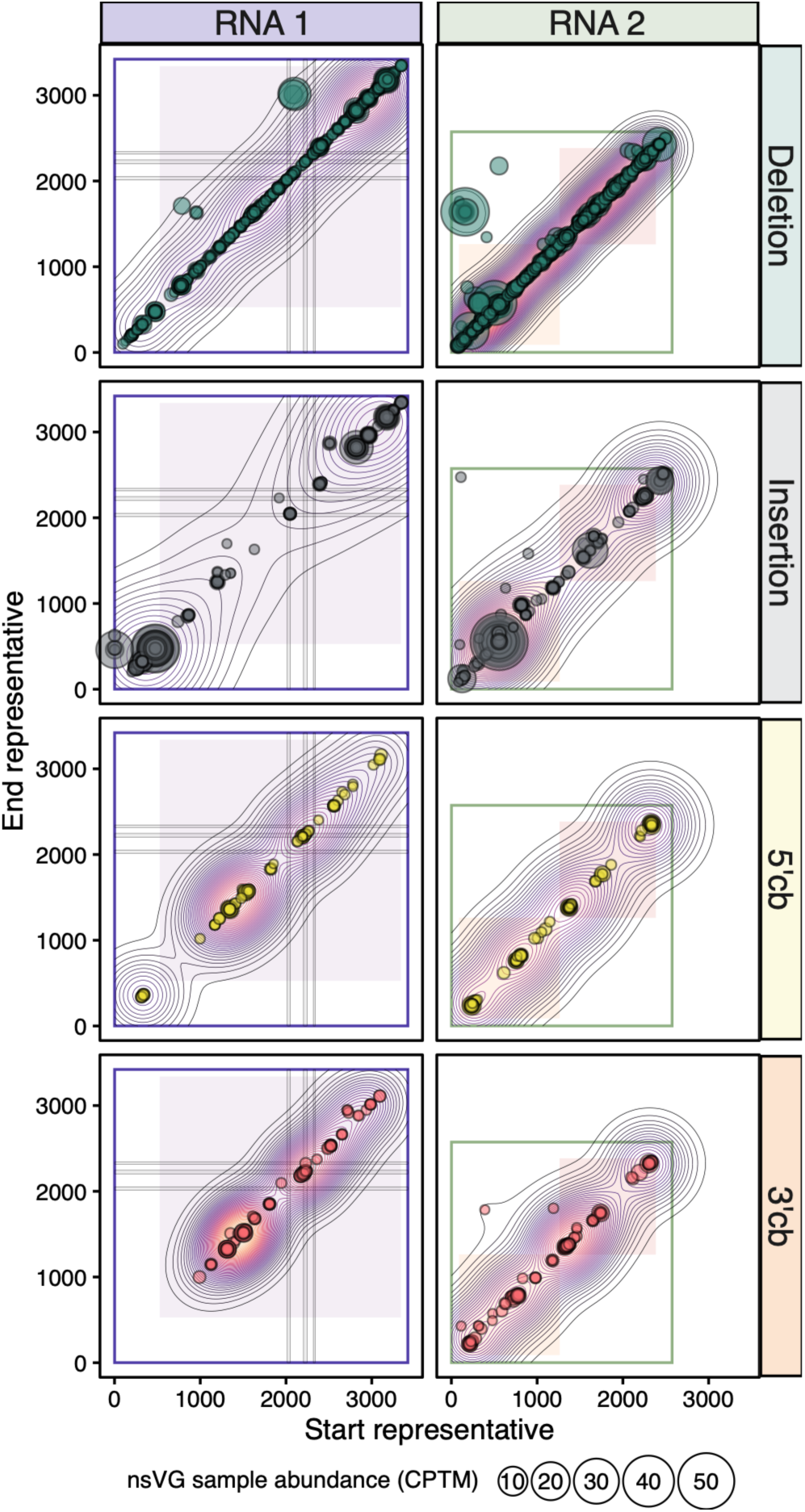
Recurrent genomic hotspots of nsVG formation. Representative start and end coordinates of clustered nsVG junctions supported by > 5 reads are shown for each nsVG category and genomic segment. Point size is proportional to cluster abundance (CPTM) at each sample, and contour lines indicate cluster density. Background shading denotes annotated coding regions, whereas gray bars indicate conserved palmprint motifs within RdRp.

Insertions exhibited a more restricted genomic distribution and accumulated in a limited number of genomic regions. Interestingly, the most recurrent insertion clusters corresponded predominantly to single-nucleotide events and ranked among the most abundant nsVG clusters identified (Fig. S7A-C). We found that in all nsVG categories, recurrence decreased significantly with event size (Spearman’s *r_S_* = −0.374, 485 d.f., *P* < 0.001; Fig. S7D), indicating that highly recurrent events generally involve short genomic modifications. Consistent with this observation, many of the most recurrent insertion and deletion clusters were located within or adjacent to homopolymeric tracts. Indeed, insertion and deletion clusters were associated with significantly more homopolymer-rich sequence contexts than cb molecules (Kruskal-Wallis test: χ^2^ = 28.938, 3 d.f., *P* < 0.001; *post hoc* Dunn test: all insertion and deletion *vs* cb comparisons adjusted *P* < 0.002; Fig. S7D), suggesting that local sequence composition contributes to the repeated generation of short nsVGs. Whether these events reflect biologically relevant polymerase slippage, technical artefacts, or a combination of both remains unclear.

Both cb types exhibited highly structured genomic distributions, with recurrent hotspots confined to a limited number of genomic regions. Although cb molecules were generally less diverse and less abundant than deletion nsVGs, the repeated detection of the same representative clusters across independent samples indicates that their formation could be likewise constrained by specific genomic features.

Despite the predominance of highly recurrent short nsVGs, one RNA2 deletion hotspot clearly differed from the general pattern. Whereas most recurrent events involved only one or a few nucleotides and were frequently associated to homopolymeric contexts (Fig. S7), this hotspot generated a deletion spanning more than 1,000 nucleotides while remaining among the most abundant deletion-derived nsVGs in the dataset (Fig. 9 and Fig. S7B). Because of its abundance, recurrence, and genomic organization, we investigated this deletion further as a candidate subgenomic RNA.

### A recurrent RNA2 deletion defines a candidate subgenomic RNA

Representative clustering identified a deletion connecting positions near the 5’ end of RNA2 with a region located within the δ sequence (representative cluster RNA2:Deletion_160_1640), which was detected in six independent samples and accumulated to substantially higher abundances than expected for deletion events of comparable size.

Inspection of the underlying junctions revealed additional closely related deletion variants mapping to the same genomic region, consistent with a recurrent fuzzy deletion hotspot spanning approximately positions 98-187 to 1531-1658 within RNA2. Similar junctions were first detected at 12 hpi and recurred throughout infection (Fig S8). Remarkably, the same deletion hotspot was also identified in independent sequencing datasets, including experimentally evolved OrV populations (Castiglioni, Olmo-Uceda, Martín et al. 2024), where it likewise represented one of the most abundant deletion nsVGs. Together, these observations indicate that this RNA2 rearrangement is repeatedly generated across independent infections and experimental contexts.

All reconstructed deletion variants in this hotspot retained an AUG at position 1672 within RNA2 (Fig. S8A-B). The AUG is in frame with the δ ORF and predicts a product lacking the N-terminal 109 amino acids of full-length δ (Fig. S8C-D).

To assess the evolutionary context of this putative alternative initiation site, we examined the corresponding region in all publicly available nematode-infecting noda-like viruses. The AUG associated with the candidate truncated δ was conserved across all available OrV isolates and was also present in LEBV. Although the equivalent methionine was not retained in the currently available MELV and SANTV sequences, the surrounding protein region remained highly conserved. Thus, while conservation alone does not imply biological function, the candidate start codon is located within a region that appears to be under evolutionary constraint.

To explore the structural consequences of initiating translation from this alternative AUG, we predicted the structure of the putative truncated δ using AlphaFold 3.0 (Fig. S8C-D) (Abramson et al. 2024). The resulting model retained the characteristic pentameric organization previously described for δ, included the C-terminal globular domain together with the elongated fiber-like region. However, the predicted protein lacked the N-terminal CP-proximal fiber segment present in the full-length. Structural alignment between the experimentally resolved δ (1-101) crystal structure (PDB 5W82; Guo and Tao 2018; Guo et al. 2020) and the AlphaFold 3.0 prediction of full-length δ showed strong agreement in the resolved N-terminal region, supporting the structural plausibility of the full-length model and highlighting the region absent from the truncated variant (Fig. S8C).

RT-PCR amplification confirmed the predicted junction in vivo (Fig. S9). These data validate the recurrent deletion molecule but do not establish its translation or its function as a *bona fide* sgRNA.

## DISCUSSION

By separately profiling (+)- and (−)-strand RNAs, we found that OrV replication generates distinct molecular populations that differ in abundance, nucleotide diversity, and mutational composition. (−)-strands contained greater diversity, whereas only a subset of their SNVs was also detected in (+)-strands. Together with the functional constraint observed among shared variants, these findings support a model in which viral diversity is progressively restricted during genome production. In parallel, analysis of structural variants identified recurrent nsVGs, including a validated RNA2 deletion that may represent a previously undescribed candidate subgenomic RNA.

Because experimental sampling was destructive, consecutive time points represent independent host populations rather than longitudinal observations of the same viral lineages. Consequently, changes in variant frequencies cannot be interpreted as direct evidence of within-host selection. Throughout this discussion, we use *filtering* to describe the observed loss or failed propagation of variants, whereas *purifying selection* is reserved for patterns that are consistent with, but do not directly demonstrate, preferential removal of deleterious variation.

### Asymmetric accumulation is consistent with template-constrained replication

OrV accumulation was strongly asymmetric and followed a biphasic temporal profile. The second accumulation wave was characterized by a marked delay in (+)RNA2 relative to the other viral RNA species, indicating that strand and segment accumulation become partially uncoupled during late infection. Similar multiphasic viral dynamics have been associated with temporally structured host responses, tissue-level spread, or compartmentalized replication in several RNA viruses (Baccam et al. 2006; Pawelek et al. 2012; Ke et al. 2022; Perdoncini Carvalho et al. 2023). The precise origin of the pattern observed here remains unresolved, but the delayed accumulation of (+)RNA2 suggests segment-specific regulation of synthesis, stability, or turnover.

The pronounced asymmetry between (+)- and (−)-strands is consistent with recent microscopy-based analyses showing that OrV replication intermediates occupy distinct intracellular environments associated with viral replication complexes and antiviral RNAi pathways (Ranganathan et al. 2026). More importantly, the saturating relationship between (−)RNA and (+)RNA abundance is inconsistent with the approximately proportional increase expected if newly synthesized templates re-entered replication with equal efficiency under a strict geometric replication model. Instead, it supports a template-constrained replication regime in which only a limited or functionally restricted subset of templates contributes disproportionately to progeny synthesis: the so-called stamping machine (SMR).

Replication mode has important evolutionary consequences because geometric replication allows mutations to pass through multiple successive copying generations, whereas SMR-like restricts the number of templates contributing to future progeny. Nevertheless, geometric replication and pure SMR are best viewed as theoretical extremes. Intermediate replication strategies have been described in bacteriophages, plant RNA viruses and large DNA viruses (Chao et al. 2002; García-Villada and Drake 2012; Martínez et al. 2011; Sardanyés et al. 2012; Boezen et al. 2022). We therefore interpret the OrV pattern as evidence of template-constrained replication with a strong SMR-like component, rather than proof of a pure replication mechanism.

### (−)-strands may constitute a transient diversity reservoir

The relationship between viral accumulation and diversity was also saturating, but the maximum diversity attained differed between strands. (−)RNA consistently accumulated more diversity than (+)RNA, indicating that replication intermediates contain a broader spectrum of sequence states than is ultimately represented within the genomic population. This result is central because it separates mutation generation from successful propagation: the diversity present in replication intermediates is not simply mirrored in progeny genomes.

Most notably, a substantial fraction of SNVs detected in (−)RNA was not recovered from (+)RNA. The proportion of shared variants increased later in infection, suggesting that much of the variation observed in replication intermediates is transient or inefficiently propagated. Not all sequence states present in (−)RNA therefore contribute detectably to the amplifying (+)-strand population. Such filtering could arise through several non-exclusive mechanisms, including selective template usage, differential RNA stability, restricted access to replication complexes, intracellular bottlenecks, or host-mediated degradation.

The minimum conditional strand-overlap observed around 12 hpi may reflect more than replication-intrinsic constraint. This time point coincides with the viral-load peak and with detectable activation of OrV-induced host responses, including RNAi- and ubiquitin-associated antiviral programs (Sterken et al. 2014; Ashe et al. 2015; Chen et al. 2017; Castiglioni, Olmo-Uceda, Villena-Giménez et al. 2024). Because viral dsRNA replication intermediates are substrates for antiviral RNAi, host sensing and degradation could contribute to the selective loss of some (−)RNA-associated variants before they reach the (+)RNA population. Comparative strand-resolved diversity analyses in RNAi-defective or immunocompromised animals would therefore help separate replication-intrinsic filters from host-imposed ones.

The non-random distribution of strand-specific variants further supports this interpretation. Regions enriched in variants restricted to (−)RNA accumulated preferentially within specific portions of the genome, particularly near the C-terminal region of the polymerase. Protein-level constraints cannot operate while mutations remain confined to non-translated replication intermediates. However, once copied into (+)RNA, variants become exposed to constraints affecting replication, translation, encapsidation, RNA stability, or protein function. Strand restriction may therefore reflect the combined action of template-level bottlenecks, host-mediated degradation, and downstream functional constraints.

Although predicted RNA structure explained only part of this variation, the association between strand-specific diversity and structural asymmetry suggests that RNA architecture may contribute to template accessibility or utilization during replication. Similar effects have been proposed for (+)RNA viruses in which local RNA structures influence polymerase progression, template selection, replication efficiency, or genome packaging (Sztuba-Solińska et al. 2011; Simon-Loriere and Holmes 2011; Perdoncini Carvalho et al. 2023). The modest strength of the association in OrV indicates that RNA structural potential is likely one component of a broader set of strand-dependent constraints, rather than the sole determinant of diversity localization.

### Sequential filtering and functional constraint

Variants detected on both strands represent the subset that successfully passed the first observable apparent within-cell filter. Their strong allele-frequency concordance indicates that propagation generally preserves relative variant abundance. However, nonsynonymous variants exhibited significantly lower strand concordance than synonymous variants, suggesting that nonsynonymous mutations are already subject to differential constraints after their generation. This pattern is consistent with the action of purifying selection on newly arising variants, a process widely documented in RNA virus populations where many nonsynonymous mutations are transient and removed before contributing to longer-term evolution (Hughes and Hughes 2007; Lauring and Andino 2010).

Recurrence patterns provide an independent line of evidence for the same process. Recurrent variants occurred above the sample-specific mutational background and were disproportionately synonymous, indicating that their repeated detection cannot be explained solely by ongoing mutational input. Rather, the same synonymous variants repeatedly emerged and persisted to detectable frequencies across independent infections, whereas nonsynonymous variants did so less often. This pattern is consistent with mutational input coupled with filtering of protein-altering variation, as expected under the dynamics of RNA virus mutant spectra (Hughes and Hughes 2007; Lauring and Andino 2010; Domingo and Perales 2019).

The sliding-window analysis localized where this filtering was most consistently observed. RdRP and CP exhibited lower *d_N_ − d_S_* values than δ overall, indicating greater depletion of nonsynonymous variation in the polymerase and capsid proteins. Although individual windows in δ occasionally reached similarly extreme values, these signals were not recurrent enough across samples to produce significant hotspots, as observed for the highlighted regions in RdRP and CP. In RdRP, the significant hotspot corresponded largely to the highly recurrent synonymous variant 2401C>U (A625), located immediately adjacent to the conserved palm domain motifs, whereas in CP they clustered within a bottom-lateral region of the capsid trimer near interfaces between neighboring subunits and facing the particle interior. Such inter-subunit interfaces are often subject to strong structural constraints because they contribute to capsid assembly and stability (Cheng and Brooks 2015). Together, these observations suggest that recurrent mutational input remains detectable in these regions but accumulates predominantly as synonymous variation, consistent with stronger constraints on protein-altering mutations

Together, strand propagation, allele-frequency concordance, variant recurrence, and localized nonsynonymous depletion all support a model in which variation that reaches the (+)-strand population remains subject to additional filtering before contributing to longer-term evolution.

### nsVGs and a possible RNA2-derived subgenomic RNA

Point mutations were not the only source of OrV diversity. Infection also generated a heterogeneous population of nsVGs whose richness, effective diversity and community structure were associated primarily with viral accumulation rather than infection time. Similar relationships have been described in influenza viruses, coronaviruses and other RNA viruses, where increasing replication activity provides additional opportunities for template switching, polymerase slippage, and incomplete genome synthesis (Saira et al. 2013; Alnaji et al. 2019; Aguilar Rangel et al. 2023).

All nsVG categories accumulated at recurrent genomic hotspots, and recurrence was strongly associated with abundance, indicating that a relatively small subset of nsVGs dominates the structural diversity of the viral population. Similar hotspot formation has been reported across diverse RNA viruses and is often linked to local sequence features, template-switching preferences, or constraints acting on the replication machinery (Alnaji et al. 2021; Hillung et al. 2024).

Most highly recurrent nsVGs corresponded to short insertion and deletion events. Recurrence decreased with event length, and many recurrent events were associated with homopolymeric sequence contexts. Such patterns are compatible with polymerase slippage or transient template misalignment during replication (Simon-Loriere and Holmes 2011), although the present data cannot distinguish biological mechanisms from technical artifacts. For this reason, short homopolymer-associated nsVGs should be interpreted cautiously.

The recurrent RNA2 deletion identified here differs markedly from the majority of recurrent nsVGs. This hotspot removes more than 1000 nucleotides while remaining among the most abundant deletion-derived molecules detected. Its pervasiveness across independent infections, recurrence in experimentally evolved populations, and validation by RT-PCR argue against a purely stochastic origin.

The organization of this deletion is compatible with a previously undescribed RNA species capable of expressing a shortened version of δ from an internal in-frame AUG. Although subgenomic RNAs are common among (+)RNA viruses (Miller and Koev 2000; Sztuba-Solińska et al. 2011), they have not been reported previously in OrV. The candidate initiation codon is conserved across all available OrV isolates and is also present in LEBV, while the surrounding protein region remains conserved across currently available nematode-infecting noda-like viruses. Nevertheless, neither the presence of the deletion nor the preservation of coding potential demonstrates regulated synthesis, translation, or biological activity. We therefore regard this molecule as a recurrent RNA2 nsVG and candidate sgRNA whose functional relevance remains to be established.

### A model of short-term filtering during OrV infection

Together, our results support a sequential model of variation and constraint during primary OrV infection (Fig. 10). Template-constrained replication generates a broader spectrum of variation in (−)RNA, but only a subset of this variation becomes detectably propagated to (+)RNA. Nonsynonymous variants encounter additional constraints after reaching the functional positive-strand population. In parallel, replication generates structural diversity through nsVG formation, although the contribution of these molecules to viral fitness remains unresolved.

**Figure 10.**
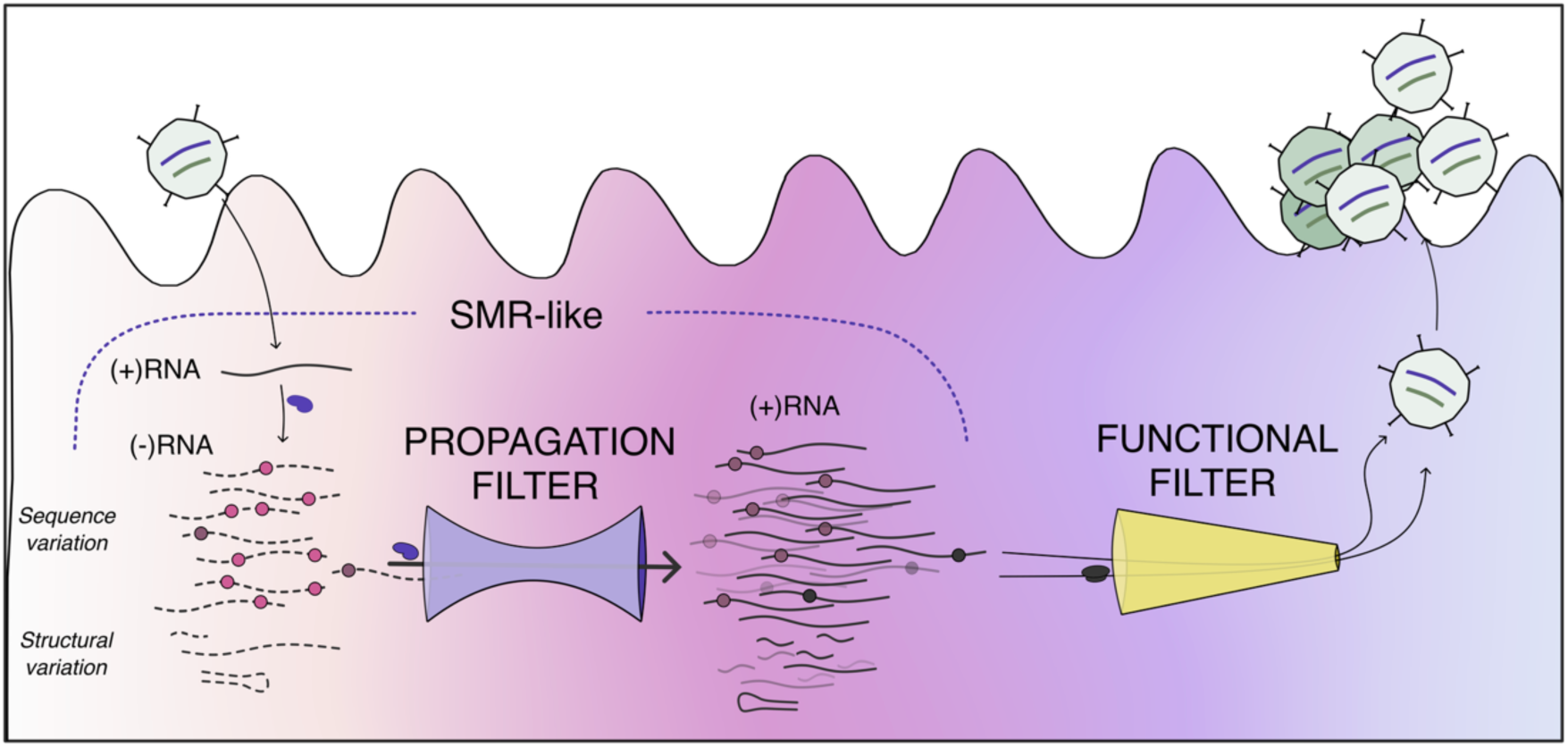
Sequential filtering model of OrV diversity during primary infection. OrV replication is represented as a template-constrained, SMR-like process in which a limited pool of (−)-strand replication intermediates gives rise to a larger population of (+)-strand progeny genomes. During replication, both sequence variation, shown as point mutations, and structural variation, shown as nsVGs, are generated within the viral population. However, only a subset of this diversity is propagated from the (−)RNA pool to the (+)RNA population, defining an early propagation filter (burgundy/brown dots). This filter may reflect the combined effects of template usage, replication-complex constraints, RNA structure, intracellular bottlenecks, and the host immune system. Variants that reach the (+)RNA population are then subjected to a second functional filter, including constraints related to RNA stability, translation, protein function, assembly, egress, and particle production. All these processes occur within a changing host-cell environment that reacts to infection and can further constrain the viral diversity that is maintained. Under this model, OrV evolutionary stability does not result from a lack of mutational input, but from the repeated restriction of newly generated variation during strand propagation and functional genome amplification.

This interpretation is consistent with broader theoretical and empirical models of within-host viral evolution, which show that observed diversity depends on the interaction between viral demography, mutation, drift, selection, bottlenecks, spatial structure, and host immune activation, rather than on mutational input alone (Fabre et al. 2012; Nelson and Hughes 2015; McCrone et al. 2018; Alamil et al. 2022). In particular, models fitted to high-throughput sequencing data have shown that selection and genetic drift jointly shape within-host viral dynamics, while stochastic bottlenecks restrict which variants persist during host colonization or transmission (Fabre et al. 2012; McCrone et al. 2018). Models integrating viral-load dynamics with mutation, drift, selection, and host immune activation further predict that within-host diversity can follow non-linear or non-monotonic trajectories rather than scaling directly with viral accumulation (Alamil et al. 2022). However, most of these frameworks treat the viral population as a single genomic compartment. Our strand-resolved data suggest an additional molecular layer of filtering, in which diversity present in replication intermediates is not necessarily transmitted to progeny genomic RNA.

Importantly, these filters are unlikely to operate independently. Viral replication takes place in a changing intracellular environment in which template availability, replication-complex organization, RNA structure, host antiviral sensing, and tissue-level physiological state can all vary over time. Under this systems-level view, the fate of a variant is not determined by replication mode, RNA sequence, protein function, or host immunity alone, but by their interaction within a temporally structured infection. The same mutation may therefore be tolerated in (−)-strand intermediate, lost during template propagation, constrained after entering the (+)-strand population, or affected by the antiviral state of the infected intestinal cell.

In this context, the minimum conditional strand-overlap observed around 12 hpi can be viewed as a transient host-virus filter, potentially arising from the overlap between replication-complex limitation and activation of antiviral pathways. This interpretation is consistent with OrV infection dynamics, in which 12 hpi coincides with the viral-load peak and with detectable host transcriptional responses, including immune- and ubiquitin-associated programs (Chen et al. 2017; Castiglioni, Olmo-Uceda, Villena-Giménez et al. 2024). It is also compatible with the central role of antiviral RNAi in OrV control, since viral dsRNA replication intermediates are substrates for DRH-1/RNAi-mediated sensing and degradation (Sterken et al. 2014; Ashe et al. 2015). Thus, strand-dependent propagation could be reflecting not only replication-intrinsic constraints, but also the changing antiviral state of infected intestinal cells.

Our results indicate that strand-resolved sequencing reveals information obscured when viral RNA is analyzed as a single population. The observed asymmetries support a model of sequential restriction of genetic diversity during OrV replication, although destructive sampling, detection thresholds, and the absence of molecular lineage tracking prevent direct reconstruction of variant transfer between RNA molecules. Future experiments combining strand-specific long-read sequencing, metabolic RNA labelling, and functional analysis of the recurrent RNA2 deletion will be required to distinguish replication, selection, degradation, and sampling effects.

## CONCLUSIONS

This study shows that OrV generates substantial short-term diversity during primary infection, but that this diversity is progressively constrained before it can contribute to the (+)-strand viral population. Strand-specific accumulation, saturating relationships between RNA abundance and diversity, and the incomplete propagation of (−)-strand variants all support a template-constrained replication regime in which (−)-strand intermediates contain a broader and more transient mutational spectrum than progeny genomic RNA.

The variants that do propagate to (+)-strands remain functionally constrained, as reflected by the predominance of synonymous recurrent mutations and localized depletion of nonsynonymous variation near structurally important regions of RdRP and CP. In addition, OrV infection generates a diverse but non-random population of nsVGs, including a recurrent RNA2 deletion compatible with a candidate subgenomic RNA encoding a truncated δ protein.

Overall, our results indicate that OrV evolutionary stability is not explained by an absence of variation. Rather, it emerges from sequential filtering acting during replication, strand propagation, host interaction, and functional genome amplification. This framework connects the intracellular replication dynamics of OrV with its limited long-term evolutionary change.

Future strand-resolved analyses in diverse host backgrounds, including animals with immunocompromised backgrounds and different developmental stages, particularly RNAi-impaired animals, will be essential to determine how much of this short-term filtering arise from intrinsic features of the OrV is due to the viral replication mode and how much is imposed by host responses.

## METHODS

### Experimental design and sample collection

Full details of the experimental design, time-course generation, sample preparation, RNA sequencing details and RNA-seq preprocessing are provided in Castiglioni, Olmo-Uceda, Villena-Giménez et al. (2024).

### Strand-specific RNA quantification

Processed FASTQ files were aligned to the OrV reference genome with BWA-MEM (Li 2013) using default parameters. Alignments were converted to BAM and sorted with SAMtools (Danecek et al. 2021). Each alignment file was then split by strand orientation. Reads mapping to the positive or negative strands were identified with bedtools bamtobed (Quinlan and Hall, 2010). For paired-end data, the following orientation rules were applied: ‘+’ of read 2 and ‘−’ of read 1 were classified as plus-strand reads, whereas ‘+’ of read 1 and ‘−’ of read 2 were classified as minus-strand reads.

Total fragment counts for each of the four viral species (+)RNA1, (+)RNA2, (−)RNA1, and (−)RNA2 (fragments*_i_*) were obtained with custom scripts from samtools coverage results on the strand-split BAMs. For each molecular species, counts were normalized by segment length (length*_i_*; (RNA1: 3421 nt and; RNA2: 2574 nt)) and library size (number of fragments in the cleaned FASTQ files) as follows: VPMB*_i_* = fragments*_i_* × 10^6^/(length*_i_* × library size). This metric assumes uniform transcription across the genome and equal sequencing accessibility for all transcripts. Coverage profiles, however, indicated this was not the case. Whether these irregularities reflect biological and/or technical noise or a more complex transcriptional process (*e.g*., the presence of nsVGs or intermediate transcripts such as subgenomic RNAs) will be addressed in detail.

### Nucleotide diversity analyses

Per-site nucleotide counts were extracted from the processed, strand-split BAM files with pysamstats v1.1.2 (Miles 2014) using the variation mode. These count tables were used to compute nucleotide diversity indices. Normalized Shannon entropy (*S_n_*) was calculated using NormShannon from QSutils v1.16.1 (Gregori et al. 2014; Gregori et al. 2016; Guerrero-Murillo and Gregori 2023). For estimates of mean normalized entropy per sample, only positions with a minimum depth of 20 reads were included.

To identify genomic positions showing strand-dependent diversity differences, nucleotide diversity values from matched (+)- and (-)-strand libraries were combined by sample, RNA segment, position, and time point. Strand bias was calculated as: Δ*S̄_n_*(*p*)=*S̄_n_*^+^(*p*)-*S̄_n_*^-^ (*p*), where positive values indicate higher diversity in the positive strand and negative values indicate higher diversity in the negative strand. Because time did not explain a major component of the position-wise residual signal, mean Δ*S̄_n_*(*p*) values were summarized by genomic position *p* and time point across biological replicates. Segment-specific empirical distributions were then used to classify positions according to the magnitude of strand bias. Positions below the 1^st^ percentile or above the 99^th^ percentile were considered extreme strand-biased positions. To identify broader genomic regions enriched in strand-biased diversity, Δ*S̄_n_*(*p*) values were smoothed using 50-nt sliding windows with 20-nt steps.

### Predicted RNA structural potential

Predicted RNA structural potential was estimated with RNAfold from the ViennaRNA package (Lorenz et al. 2011). For each RNA segment, 50-nt windows were generated every 5-nt from both the sense sequence and the reverse-complement sequence. RNAfold was run at 20 °C with partition-function calculation, isolated base pairs disallowed, and G-quadruplex formation allowed: RNAfold --noPS -p -g -T 20 –noLP.

Thus, predicted minimum free energy (MFE) values should be interpreted as reflecting both canonical RNA secondary structures and G-quadruplex-compatible conformations. For visualization, –MFE values were reported, so that larger values corresponded to higher predicted structural stability. Strand-specific structural asymmetry was calculated for matched windows as: Δ*MFE*(*p*)=*MFE*^(+)^(*p*)-*MFE*^(-)^(*p*), where negative values indicate greater predicted structural stability in the (−)-strand orientation. Windows with absolute structural asymmetry above the segment-specific 95^th^ percentile were classified as structural hotspots.

Associations between strand-dependent nucleotide diversity residuals and structural asymmetry were assessed using Spearman’s correlations. A permutation test was performed by randomly permuting absolute diversity-bias values across windows while preserving the observed structural-bias values. This procedure was repeated 5,000 times, and the empirical *P* value was calculated as the proportion of permuted correlations with absolute values greater than or equal to the observed correlation. Linear and generalized additive models including GC content were also used to assess whether the association between diversity bias and structural asymmetry persisted after accounting for local nucleotide composition.

### SNV detection and classification

Single-nucleotide variants (SNVs) were called independently from (+)- and (−)-strands alignments using LoFreq (Wilm et al. 2012), after removal of optical duplicates. Separate VCF files were generated for (+)- and (−)-strand libraries and then combined into a single table. For each SNV, genomic segment, sample, time point, replicate, strand, genomic position, reference allele, alternative allele, read depth, and allele frequency (AF) were retained.

Variants were annotated according to genomic region and coding effect. Coding SNVs were classified as synonymous or nonsynonymous by comparing the reference codon with the codon generated by the alternative nucleotide. SNV identifiers were generated from genomic segment, genomic position, and nucleotide change, with thymidine converted to uridine for reporting. Only SNVs with AF > 0.01, total depth ≥ 20, and SNV Phred quality ≥ 20 were retained for downstream analyses.

SNV richness was defined as the number of SNVs with AF > 0.01 in each sample, RNA segment, and strand. SNV abundance was defined as the cumulative AF of all SNVs above this threshold.

For strand-resolved analyses, SNVs were classified within each sample as (+)RNA-specific when detected only in (+)-strand libraries, (−)RNA-specific when detected only in (−)-strand libraries, and shared when detected in both strands. For each unique SNV, recurrence was quantified as the number of independent samples in which the variant was detected above the AF threshold.

### Cross-strand SNV overlap

To estimate the conditional strand overlap for variants detected in (−)-strand replication intermediates were also detected in (+)-strand RNA, we calculated, for each sample and RNA segment: *P*(shared | −) = *N*_shared_/(*N*_shared_ + *N*(−)_specific_); where *N*_shared_ is the number of SNVs detected on both strands and *N*(−)_specific_ is the number of SNVs detected only on the (−)-strand. Temporal variation in this conditional strand-overlap was analyzed separately for RNA1 and RNA2 using quasi-binomial generalized linear models of the form: cbind(transmission_success, transmission_total − transmission_success) ∼ factor(hpi); where transmission_success corresponds to the number of shared SNVs and transmission_total corresponds to all SNVs detected on the negative strand. Pairwise comparisons among time points were performed on the log-odds scale using estimated marginal means with BH correction.

### Genomic enrichment and recurrence of SNVs

To assess whether strand-specific SNVs were randomly distributed along each genomic segment, KDEs were calculated separately for (+)RNA-specific and (−)RNA-specific SNVs in RNA1 and RNA2. Each SNV was weighted by its recurrence across independent samples. KDEs were calculated using a bandwidth of 100 nt and evaluated over 512 equally spaced genomic positions.

Statistical significance was evaluated using a permutation-based null model. For each RNA segment and strand-specific category, SNV positions were randomly reassigned across the corresponding genomic segment while preserving both the number of SNVs and their recurrence weights. This procedure was repeated 5,000 times. Pointwise 95% confidence envelopes were calculated from the permuted KDEs, and genomic regions where the observed KDE exceeded the upper confidence limit were considered enriched for strand-specific SNVs beyond random expectation.

### AF concordance between strands

For SNVs detected on both strands within the same sample, AF concordance was assessed by comparing AF values measured in (−)RNA and (+)RNA libraries. Only SNVs detected with AF > 0.01 on both strands were included. Linear models were used to test whether the relationship between AF in (−)RNA and AF in (+)RNA differed according to predicted coding effect or genomic region. In these models, AF in (+)RNA was used as the response variable and AF in (−)RNA as the predictor. Interaction terms were used to test whether slopes differed between synonymous and nonsynonymous variants or among viral proteins.

Recurrent shared SNVs were summarized by counting the number of independent samples in which each SNV was detected on both strands above the AF threshold. For visualization of recurrent shared variants, SNVs detected on both strands in at least four samples were displayed as a heatmap using AF values measured in (+)-strand libraries.

### Analysis of selective constraints

To characterize local selective constraints across viral coding regions, we calculated *d_N_* − *d_S_* in overlapping four-codon sliding windows with a one-codon step. Because this analysis focuses on codon effects, only coding SNVs detected in (+)RNA with AF > 0.01 were included.

For each codon, the expected numbers of synonymous and nonsynonymous sites were calculated using the Nei and Gojobori (1986) method. Within each window and sample, observed synonymous and nonsynonymous mutations were summed and normalized by the corresponding numbers of synonymous and nonsynonymous sites to obtain *p_S_* and *p_N_*. Jukes and Cantor (1969) correction was then applied to transform these values into *d_S_* and *d_N_*, respectively. Windows for which the correction was undefined were treated as missing values. Global differences in *d_N_* − *d_S_* among viral proteins were assessed using Kruskal-Wallis tests followed by Bonferroni-Hochberg corrected pairwise Wilcoxon rank-sum tests. Temporal changes in window-level *d_N_* − *d_S_* were analyzed using linear mixed-effects models including protein and time as fixed effects and sample identity as a random effect. The protein × time interaction was evaluated by model comparison but was not retained when it did not improve model fit.

To identify local windows showing significant deviations from neutrality, one-sided Wilcoxon signed-rank tests were applied to each window across samples against the null expectation *d_N_* − *d_S_* = 0. Tests were performed independently for positive deviations (*d_N_* − *d_S_* > 0) and negative deviations (*d_N_* − *d_S_* < 0). Bonferroni-Hochberg correction was applied separately within each protein. Windows with adjusted *P* < 0.05 were considered significantly enriched for nonsynonymous or synonymous variation, respectively.

### nsVG detection, clustering and quantification

NsVGs were identified and characterized with DVGfinder (Olmo-Uceda et al. 2022). We used the metasearch mode, but only nsVGs detected by ViReMa were retained for downstream analyses (Routh and Johnson 2014). Because strand information was ignored for this section, the two junction coordinates were referred to as start, the junction closest to the 5’ end, and the junction closest to the 3’ end.

Unique nsVG junctions were defined by RNA segment, nsVG category, and start/end coordinates. To account for minor variability in junction mapping, junctions were clustered independently for each RNA segment and nsVG category using a representative-centered spatial clustering strategy. Junctions were first ranked according to their cumulative abundance across all samples, using CPTM followed by read count. The most abundant unassigned junction was selected as the cluster representative, and all remaining junctions located within ±10 nt of both its start and end coordinates were assigned to the same cluster. This process was repeated iteratively until all junctions were assigned.

This approach groups nearby junctions while preventing the chaining effect that can occur with conventional density-based clustering methods, whereby distant junctions become connected through a series of intermediate positions. Cluster abundances were subsequently calculated by summing all reads assigned to the corresponding representative cluster within each sample. Only representative clusters supported by more than five reads in a sample were retained for downstream analyses.

NsVG richness was defined as the number of distinct representative clusters detected in each sample, RNA segment, and nsVG category. NsVG abundance was quantified in two ways. First, counts per million mapped reads (CPTM) were used for community-level diversity analyses because they normalize for library size. Second, relative viral abundance was calculated as: CPVT = 1000 × nsVG-supporting reads/viral reads from the corresponding RNA segment. This measure expresses nsVG abundance as nsVG-supporting reads per 1000 viral reads and was used when evaluating nsVG accumulation relative to the viral population.

NsVG richness and relative abundance were modelled as functions of viral accumulation, RNA segment, and nsVG category using generalized mixed-effects models. The contribution of infection time was evaluated by comparing models with and without hpi after accounting for VPMB. Relationships between nsVG richness and relative abundance were evaluated using generalized additive models (GAMs).

### Alpha and beta diversity of nsVG communities

For community-level analyses, a sample-by-cluster abundance matrix was constructed using representative cluster abundances expressed as CPTM. Alpha diversity was quantified using richness, Shannon diversity, effective Shannon diversity, and Pielou’s evenness. Effective Shannon diversity was calculated as exp(*H*’), where *H*’ is Shannon diversity, and represents the effective number of equally abundant nsVG clusters. Pielou’s evenness was calculated as *H*’/log*S*, where *S* is richness, and was calculated only for samples with richness greater than one.

Beta diversity was assessed using Bray-Curtis dissimilarities calculated from the CPTM abundance matrix. Associations between community composition, viral accumulation, and hpi were tested using PERMANOVA with 9,999 permutations. Homogeneity of multivariate dispersion among time points was assessed using betadisper and permutation tests. These analyses were performed with vegan (Oksanen et al., 2026).

### NsVG recurrence, genomic hotspots and homopolymeric context

For each representative nsVG cluster, recurrence was defined as the number of independent samples in which the cluster was detected after abundance filtering. Total cluster abundance was calculated as the sum of CPTM values across all samples. Differences in recurrence among RNA segments and nsVG categories were analyzed using negative-binomial generalized linear models including RNA segment, nsVG category, and their interaction as predictors. A Poisson model was initially evaluated but rejected because of overdispersion. Associations between recurrence, total abundance, event size, and homopolymeric context were assessed using Spearman’s correlations.

Event size was defined as the absolute distance between representative start and end coordinates: |Δ| = |end − start|

To assess local homopolymeric context, each genomic position was assigned the length of the homopolymer tract containing that position. For example, all positions within a run of 11 consecutive cytosines were assigned a value of 11. Representative clusters were annotated with the homopolymer length at both start and end coordinates, and the maximum of these two values was used as a summary measure of local homopolymeric context. Differences in homopolymeric context among nsVG categories were assessed using Kruskal-Wallis tests followed by Bonferroni-Hochberg adjusted Dunn *post hoc* tests using the dunn.test package (Dinno 2026).

### Protein structure prediction for the recurrent RNA2 deletion

The structure of the complete δ protein and the putative truncated δ protein was predicted using AlphaFold 3.0 (Abramson et al. 2024). The experimentally resolved OrV δ N-terminal fragment, residues 1-101, was obtained from the Protein Data Bank (PDB ID: 5W82; Guo and Tao, 2018). Structural visualization and alignment were performed in UCSF ChimeraX v1.8 using Matchmaker (Pettersen et al. 2021).

### Identification and validation of the recurrent RNA2 deletion

Populations of ∼500 control and inoculated animals were prepared, collected, and total RNA was extracted as previously described (Castiglioni, Olmo-Uceda, Villena-Giménez et al. 2024). Reverse transcription was performed from 20 ng of total RNA using AccuScript High-Fidelity Reverse Transcriptase (Agilent Technologies) and the reverse primer (5’-GGGTACTCTCGGCAATCTCG). The resulting cDNA was amplified using Phusion High-Fidelity DNA Polymerase (Thermo Fisher Scientific) with the forward (5’-CGACCCGTAAAGACCTCGAC) and reverse primers. PCR products were electrophoresed on a 1.5% agarose gel with NZYDNA ladder VII (NZYtech) and GreenSafe Premium (NZYtech). A plasmid containing the full sequence of OrV RNA2 in a pBSK backbone was used as a positive control in the PCR.

### Statistical analysis

All statistical analyses were performed in R v4.2.1 (R Core Team 2022) within RStudio 2022.2.3.492. Data manipulation and visualization were performed using the tidyverse framework (Wickham et al. 2019). Linear and generalized linear mixed-effects models were fitted using lme4 (Bates et al. 2015), which implements maximum-likelihood and restricted maximum-likelihood estimation for mixed models. Estimated marginal means and pairwise contrasts were obtained using emmeans (Lenth 2026). Generalized additive models were fitted using mgcv (Wood 2017), which implements generalized additive models with automatic smoothness estimation. Community ecology analyses were performed using vegan (Oksanen et al. 2026). Dynamic time warping (DTW) analyses were performed using the dtw package (Giorgino 2009).

Model-specific distributions, link functions, fixed effects, random effects and *post hoc* tests are described for each analysis. When appropriate, likelihood-ratio tests were used to compare nested models, and multiple testing correction was performed using BH adjustment.

## Ethicas Approval

Not applicable.

## Data availability

RNA-seq data can be downloaded from NCBI SRA: PRJNA1037048. R and bash codes for sequence data analysis and presentation are available in https://github.com/MJmaolu/short-term-OrV-accumulation-and-diversification/tree/main

## Author contributions

MJOU and SFE conceptualized the idea of the study. MJOU curated the data. DH performed experimental validation. VGC generated the data. MJOU wrote the code and analyzed the data. MJOU and SFE performed formal statistical analysis. MJOU generated visualization. SFE acquired funding. MJOU and SFE wrote the manuscript. DH and VGC revised and edited the manuscript.

## Acknowledgements

We thank Francisca de la Iglesia for excellent technical support and the members of the EvolSysVir lab for their insightful comments. We especially thank Josep Sardanyés for drawing our attention to strand asymmetry and for motivating the development of a strand-sensitive approach. Computations were performed on the HPC cluster Garnatxa at I^2^SysBio.

