## Supplementary materials for "The hidden diversity: Replication-strand asymmetry shapes short-term nucleotide and structural variation in an RNA virus"

**Supplementary Text 1. Characterization of the nsVG community**

To further characterize the structure of nsVG communities, we calculated effective Shannon diversity, Pielou's evenness and Bray-Curtis dissimilarities among samples (Fig. S6). Effective Shannon diversity increased strongly with viral accumulation (linear model, $R^2 = 0.89$, $P < 0.001$; Fig. S8B), indicating that the effective number of abundant nsVG clusters increased as viral replication progressed. Consistent with the richness analyses presented in the main text, inclusion of hpi did not improve model fit after accounting for VPMB (ANOVA, $P = 0.732$).

In contrast, Pielou's evenness exhibited a weak but significant negative association with viral accumulation (effect = –0.009 ±0.004, $P = 0.041$; Fig. S5D), suggesting that increasing diversity was accompanied by a moderate increase in dominance by a subset of nsVG clusters. Again, hpi did not explain additional variation beyond VPMB (ANOVA, $P = 0.972$). Together, these results indicate that increasing viral accumulation promotes the generation of a larger and more diverse nsVG community, while simultaneously favoring the disproportionate expansion of a subset of highly represented nsVG clusters.

Analyses of beta diversity yielded similar conclusions. Bray-Curtis dissimilarities revealed that viral accumulation was significantly associated with variation in nsVG community structure (PERMANOVA, $R^2 = 0.274$, $P < 0.001$), whereas infection time was not ($R^2 = 0.045$, $P = 0.184$). The same pattern was observed when both variables were included in the model simultaneously, with VPMB remaining a significant predictor of community structure ($P < 0.001$), whereas hpi did not ($P$ = 0.184). These results are consistent with the alpha-diversity analyses and further support viral accumulation, rather than infection time itself, as the primary factor associated with differences among nsVG communities. No evidence of heterogeneous multivariate dispersion among time points was detected (PERMDISP, $P$ = 0.400), indicating that the observed differences were not driven by unequal within-group variation.

**Table S1.** Results of the Pearson's $\chi^2$ test for count data performed between the consecutive passages and between each passage and the proportions at the origin (0 hpi). Adjusted *P* values were obtained with the Benjamini-Hochberg method.

| Comparison (hpi) | $\chi^2$ | Adjusted *P* | Comparison (hpi) | $\chi^2$ | Adjusted *P* |
|---|---|---|---|---|---|
| 0 - 2 | 28.94 | 0.016 | 0 - 2 | 28.94 | 0.016 |
| 2 - 4 | 31.08 | 0.009 | 0 - 4 | 52.12 | < 0.001 |
| 6 - 8 | 5980.57 | < 0.001 | 0 - 8 | 4323.07 | < 0.001 |
| 8 - 12 | 42451.91 | < 0.001 | 0 - 12 | 6432.98 | < 0.001 |
| 12 - 18 | 45062.99 | < 0.001 | 0 - 18 | 7642.31 | < 0.001 |
| 18 - 22 | 18539.37 | < 0.001 | 0 - 22 | 9415.49 | < 0.001 |
| 22 - 26 | 66668.69 | < 0.001 | 0 - 26 | 3622.79 | < 0.001 |
| 26 - 32 | 15879.98 | < 0.001 | 0 - 32 | 8395.40 | < 0.001 |
| 32 - 38 | 12399.04 | < 0.001 | 0 - 38 | 1477.74 | < 0.001 |
| 38 - 44 | 13477.51 | < 0.001 | 0 - 44 | 11936.46 | < 0.001 |

**Table S2**. Model comparison for (+)RNA accumulation as a function of (−)RNA abundance. Nonlinear saturation and exponential (log-linear) models were fitted independently for RNA1 and RNA2. Model performance is summarized using Akaike's Information Criterion (AIC) and residual sum of squares (RSS). For each RNA segment, ΔAIC is calculated relative to the best-supported model (ΔAIC = 0). Lower AIC and RSS values indicate improved fit. The saturation model is defined as (for $i$ = 1, 2): $(+)RNA_i = a_i(-)RNA_i/[b_i + (-)RNA_i]$. The saturating model was overwhelmingly supported over the exponential model for both RNA segments (ΔAIC = 167 for RNA1; ΔAIC = 111 for RNA2). Corresponding Akaike weights strongly favored the saturating model (AICw ≈ 1.00 for both RNA segments), with negligible support for the exponential model (AICw ≈ 0 for both RNA segments), indicating essentially no evidence for exponential scaling.

| **RNA** | **Model** | **AIC** | **RSS** | **ΔAIC** | **AICw** |
|---|---|---|---|---|---|
| RNA1 | Saturation | −11.3 | 1.3 | 0 | 1 |
| | Exponential | 155.0 | 1545.3 | 167 | $10^{-37}$ |
| RNA2 | Saturation | 55.1 | 8.2 | 0 | 1 |
| | Exponential | 165.6 | 33422.2 | 110 | $10^{-25}$ |

**Table S3**. Bootstrap-based comparisons of saturation model parameters between RNA segments and strand polarities. For each comparison, differences in parameters ($\Delta a$ or $\Delta b$) were calculated from 1000 bootstrap resamplings of the original datasets, with nonlinear saturation models refitted at each iteration. Reported values correspond to the mean parameter difference and the associated 95% bootstrap confidence interval. Significant differences bolded.

| **Comparison** | **Parameter** | **Mean Δ [95% CI]** | ***P*** |
|---|---|---|---|
| **(-)RNA 1 *vs* (+)RNA 1** | ***a*** | **0.0066 [0.0053, 0.0080]** | **< 0.001** |
| **(-)RNA 2 *vs* (+)RNA 2** | ***a*** | **0.0035 [0.0027, 0.0044]** | **< 0.001** |
| **(-)RNA 1 *vs* (-)RNA 2** | ***a*** | **0.0039 [0.0025, 0.0053]** | **< 0.001** |
| (+)RNA 1 *vs* (+)RNA 2 | *a* | 0.0007 [–0.0005, 0.0014] | 0.086 |
| (-)RNA 1 *vs* (+)RNA 1 | *b* | 0.0014 [–0.0044, 0.0178] | 0.680 |
| (-)RNA 2 *vs* (+)RNA 2 | *b* | 0.0002 [–0.0049, 0.0069] | 0.714 |
| (-)RNA 1 *vs* (-)RNA 2 | *b* | 0.0014 [–0.0033, 0.0058] | 0.554 |
| (+)RNA 1 *vs* (+)RNA 2 | *b* | 0.0001 [–0.0165, 0.0083] | 0.824 |

**Table S4**. Comparison of allele frequencies (AF) between recurrent (R) and background (B) SNVs for each sample and recurrence threshold. Recurrent SNVs were defined as variants detected in more than one (> 1), two (> 2), or three (> 3) samples. Differences in AF distributions were assessed using a one-sided Wilcoxon rank-sum test. The effect size was calculated as the difference between the median AF of recurrent and background SNVs. *P* values were adjusted for multiple testing using the Benjamini-Hochberg. Significant (adjusted $P < 0.05$) are highlighted in yellow.

| | | Recurrent > 1 | | | | | | | Recurrent > 2 | | | | | | | Recurrent > 3 | | | | | | |
|---|---|---|---|---|---|---|---|---|---|---|---|---|---|---|---|---|---|---|---|---|---|---|
| **Sample** | **hpi** | **R** | **B** | **median R** | **median B** | **Effect size** | ***p*** | **FDR** | **R** | **B** | **median R** | **median B** | **Effect size** | ***p*** | **FDR** | **R** | **B** | **median R** | **median B** | **Effect size** | ***p*** | **FDR** |
| orv_2h_2 | 2 | 0 | 1 | NA | 0,0833 | NA | NA | NA | 0 | 1 | NA | 0,0833 | NA | NA | NA | 0 | 1 | NA | 0,0833 | NA | NA | NA |
| orv_4h_1 | 4 | 0 | 1 | NA | 0,0909 | NA | NA | NA | 0 | 1 | NA | 0,0909 | NA | NA | NA | 0 | 1 | NA | 0,0909 | NA | NA | NA |
| orv_4h_2 | 4 | 1 | 3 | 0,1667 | 0,0769 | 0,0897 | 0,1855 | 0,2460 | 1 | 3 | 0,1667 | 0,0769 | 0,0897 | 0,1855 | 0,2505 | 1 | 3 | 0,1667 | 0,0769 | 0,0897 | 0,1855 | 0,2505 |
| orv_6h_1 | 6 | 2 | 7 | 0,0767 | 0,0625 | 0,0142 | 0,7679 | 0,7974 | 1 | 8 | 0,1373 | 0,0502 | 0,0871 | 0,2806 | 0,3608 | 1 | 8 | 0,1373 | 0,0502 | 0,0871 | 0,2806 | 0,3608 |
| orv_6h_2 | 6 | 1 | 3 | 0,0190 | 0,1500 | - 0,1310 | 0,9632 | 0,9632 | 1 | 3 | 0,0190 | 0,1500 | - 0,1310 | 0,9632 | 0,9632 | 1 | 3 | 0,0190 | 0,1500 | - 0,1310 | 0,9632 | 0,9632 |
| orv_6h_3 | 6 | 2 | 5 | 0,1044 | 0,0556 | 0,0489 | 0,1665 | 0,2460 | 2 | 5 | 0,1044 | 0,0556 | 0,0489 | 0,1665 | 0,2365 | 2 | 5 | 0,1044 | 0,0556 | 0,0489 | 0,1665 | 0,2365 |
| orv_8h_1 | 8 | 12 | 52 | 0,0336 | 0,0043 | 0,0293 | 0,0005 | 0,0021 | 8 | 56 | 0,0425 | 0,0045 | 0,0381 | 0,0002 | 0,0013 | 7 | 57 | 0,0486 | 0,0045 | 0,0441 | 0,0005 | 0,0048 |
| orv_8h_2 | 8 | 3 | 14 | 0,0378 | 0,0312 | 0,0066 | 0,3296 | 0,4045 | 3 | 14 | 0,0378 | 0,0312 | 0,0066 | 0,3296 | 0,4045 | 3 | 14 | 0,0378 | 0,0312 | 0,0066 | 0,3296 | 0,4045 |
| orv_8h_3 | 8 | 15 | 34 | 0,0219 | 0,0043 | 0,0176 | 0,0016 | 0,0063 | 11 | 38 | 0,0219 | 0,0049 | 0,0169 | 0,0043 | 0,0147 | 8 | 41 | 0,0526 | 0,0051 | 0,0475 | 0,0026 | 0,0077 |
| orv_12h_1 | 12 | 17 | 91 | 0,0110 | 0,0025 | 0,0085 | 0,0002 | 0,0011 | 13 | 95 | 0,0134 | 0,0025 | 0,0109 | 0,0001 | 0,0007 | 9 | 99 | 0,0347 | 0,0027 | 0,0320 | 0,0009 | 0,0060 |
| orv_12h_2 | 12 | 23 | 136 | 0,0059 | 0,0023 | 0,0036 | 0,0001 | 0,0011 | 13 | 146 | 0,0113 | 0,0023 | 0,0089 | 0,0000 | 0,0001 | 9 | 150 | 0,0265 | 0,0023 | 0,0242 | 0,0000 | 0,0000 |
| orv_12h_3 | 12 | 25 | 107 | 0,0067 | 0,0028 | 0,0039 | 0,0002 | 0,0011 | 15 | 117 | 0,0123 | 0,0030 | 0,0094 | 0,0001 | 0,0008 | 9 | 123 | 0,0290 | 0,0030 | 0,0259 | 0,0001 | 0,0011 |
| orv_18h_1 | 18 | 8 | 30 | 0,0433 | 0,0160 | 0,0273 | 0,0058 | 0,0169 | 6 | 32 | 0,0433 | 0,0168 | 0,0265 | 0,0261 | 0,0470 | 6 | 32 | 0,0433 | 0,0168 | 0,0265 | 0,0261 | 0,0503 |
| orv_18h_2 | 18 | 12 | 47 | 0,0484 | 0,0191 | 0,0292 | 0,0001 | 0,0011 | 10 | 49 | 0,0463 | 0,0199 | 0,0264 | 0,0015 | 0,0069 | 8 | 51 | 0,0493 | 0,0202 | 0,0291 | 0,0013 | 0,0061 |
| orv_18h_3 | 18 | 13 | 41 | 0,0404 | 0,0261 | 0,0143 | 0,0286 | 0,0515 | 8 | 46 | 0,0627 | 0,0266 | 0,0361 | 0,0250 | 0,0470 | 7 | 47 | 0,0552 | 0,0270 | 0,0282 | 0,0783 | 0,1244 |
| orv_22h_1 | 22 | 9 | 47 | 0,0289 | 0,0073 | 0,0216 | 0,0136 | 0,0282 | 7 | 49 | 0,0289 | 0,0073 | 0,0216 | 0,0106 | 0,0260 | 6 | 50 | 0,0375 | 0,0072 | 0,0303 | 0,0011 | 0,0060 |
| orv_22h_2 | 22 | 17 | 49 | 0,0154 | 0,0076 | 0,0078 | 0,0636 | 0,1011 | 10 | 56 | 0,0220 | 0,0076 | 0,0143 | 0,0296 | 0,0499 | 8 | 58 | 0,0327 | 0,0075 | 0,0252 | 0,0026 | 0,0077 |
| orv_22h_3 | 22 | 17 | 59 | 0,0231 | 0,0065 | 0,0167 | 0,0000 | 0,0001 | 11 | 65 | 0,0187 | 0,0069 | 0,0118 | 0,0003 | 0,0015 | 8 | 68 | 0,0296 | 0,0076 | 0,0220 | 0,0022 | 0,0077 |
| orv_26h_1 | 26 | 19 | 74 | 0,0144 | 0,0057 | 0,0087 | 0,0071 | 0,0174 | 13 | 80 | 0,0144 | 0,0061 | 0,0083 | 0,0128 | 0,0288 | 9 | 84 | 0,0179 | 0,0061 | 0,0118 | 0,0044 | 0,0119 |
| orv_26h_2 | 26 | 14 | 65 | 0,0164 | 0,0047 | 0,0116 | 0,0063 | 0,0169 | 10 | 69 | 0,0174 | 0,0049 | 0,0124 | 0,0053 | 0,0158 | 8 | 71 | 0,0352 | 0,0049 | 0,0303 | 0,0050 | 0,0123 |
| orv_26h_3 | 26 | 13 | 36 | 0,0238 | 0,0103 | 0,0135 | 0,0264 | 0,0510 | 7 | 42 | 0,0123 | 0,0118 | 0,0005 | 0,1621 | 0,2365 | 6 | 43 | 0,0314 | 0,0112 | 0,0203 | 0,0424 | 0,0716 |
| orv_32h_1 | 32 | 8 | 43 | 0,0424 | 0,0097 | 0,0327 | 0,0125 | 0,0282 | 6 | 45 | 0,0669 | 0,0097 | 0,0573 | 0,0037 | 0,0144 | 5 | 46 | 0,0565 | 0,0103 | 0,0462 | 0,0163 | 0,0338 |
| orv_32h_2 | 32 | 11 | 42 | 0,0481 | 0,0083 | 0,0398 | 0,0053 | 0,0169 | 6 | 47 | 0,0529 | 0,0108 | 0,0421 | 0,0076 | 0,0205 | 6 | 47 | 0,0529 | 0,0108 | 0,0421 | 0,0076 | 0,0171 |
| orv_32h_3 | 32 | 15 | 60 | 0,0135 | 0,0086 | 0,0049 | 0,0314 | 0,0529 | 13 | 62 | 0,0135 | 0,0086 | 0,0049 | 0,0254 | 0,0470 | 9 | 66 | 0,0157 | 0,0087 | 0,0070 | 0,0288 | 0,0518 |
| orv_38h_1 | 38 | 4 | 8 | 0,0960 | 0,0584 | 0,0377 | 0,1751 | 0,2460 | 2 | 10 | 0,1818 | 0,0632 | 0,1186 | 0,3736 | 0,4386 | 2 | 10 | 0,1818 | 0,0632 | 0,1186 | 0,3736 | 0,4386 |
| orv_38h_2 | 38 | 6 | 5 | 0,0955 | 0,0349 | 0,0606 | 0,3921 | 0,4603 | 6 | 5 | 0,0955 | 0,0349 | 0,0606 | 0,3921 | 0,4411 | 4 | 7 | 0,0327 | 0,0588 | - 0,0261 | 0,8025 | 0,8333 |
| orv_38h_3 | 38 | 3 | 3 | 0,1166 | 0,0417 | 0,0750 | 0,1914 | 0,2460 | 1 | 5 | 0,5046 | 0,0667 | 0,4379 | 0,1208 | 0,1918 | 1 | 5 | 0,5046 | 0,0667 | 0,4379 | 0,1208 | 0,1812 |
| orv_44h_1 | 44 | 0 | 9 | NA | 0,1153 | NA | NA | NA | 0 | 9 | NA | 0,1153 | NA | NA | NA | 0 | 9 | NA | 0,1153 | NA | NA | NA |
| orv_44h_2 | 44 | 3 | 9 | 0,0918 | 0,0441 | 0,0477 | 0,4267 | 0,4800 | 2 | 10 | 0,0508 | 0,0612 | - 0,0104 | 0,7044 | 0,7315 | 2 | 10 | 0,0508 | 0,0612 | - 0,0104 | 0,7044 | 0,7607 |
| orv_44h_3 | 44 | 1 | 4 | 0,1969 | 0,2216 | - 0,0247 | 0,6382 | 0,6892 | 1 | 4 | 0,1969 | 0,2216 | - 0,0247 | 0,6382 | 0,6892 | 1 | 4 | 0,1969 | 0,2216 | - 0,0247 | 0,6382 | 0,7179 |

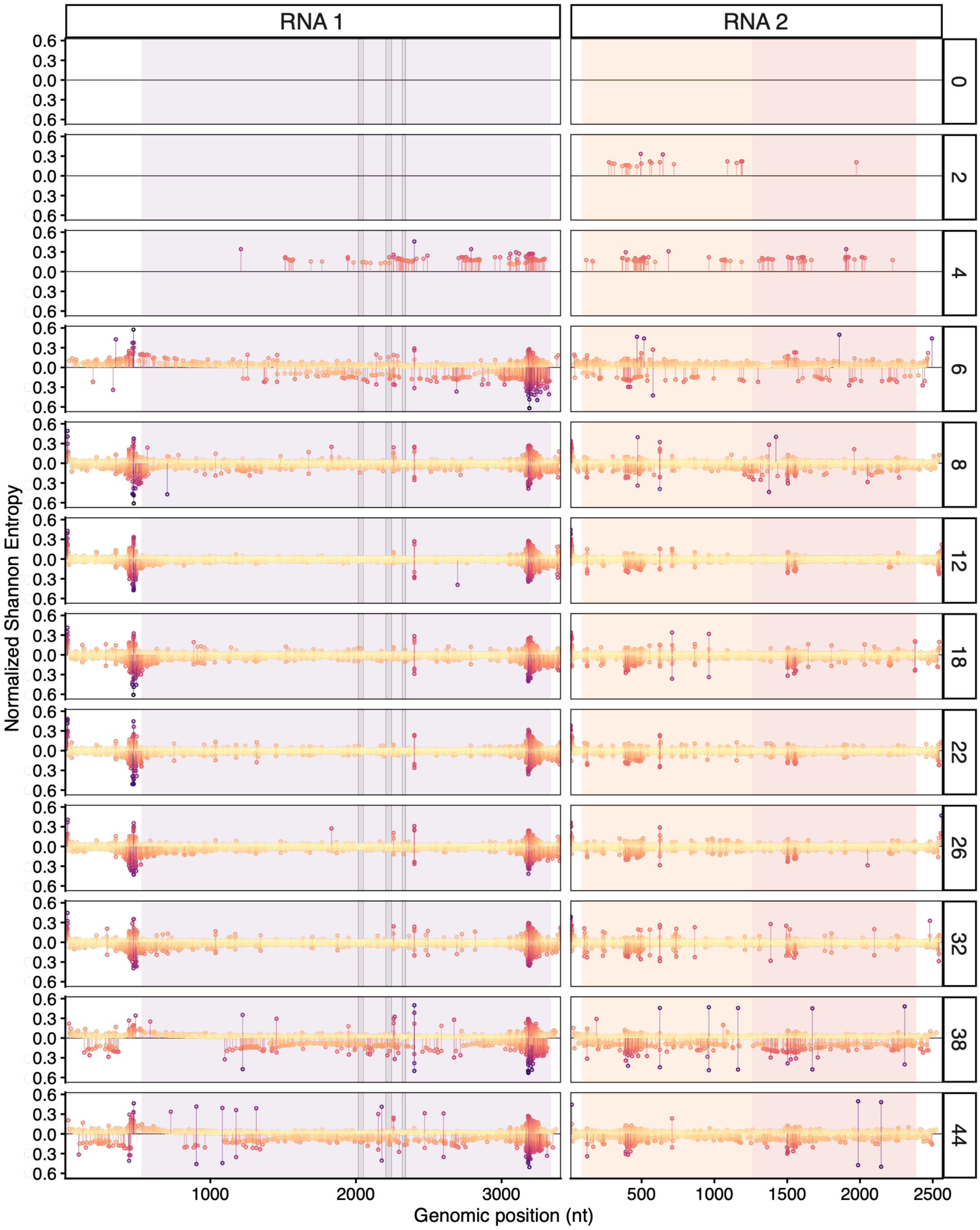


**Figure S1.** Distribution of $S_n$ along the viral genome. Per site $S_n$ was calculated independently for each strand; only positions with > 10 reads were included. For clarity, sites with $S_n = 0$ are uncolored. Dot and segment colors range from low entropy (yellow) to high entropy (purple). Background shading indicates coding regions. Gray rectangles mark the A, B, and C motifs of the RdRP palmprint domain.

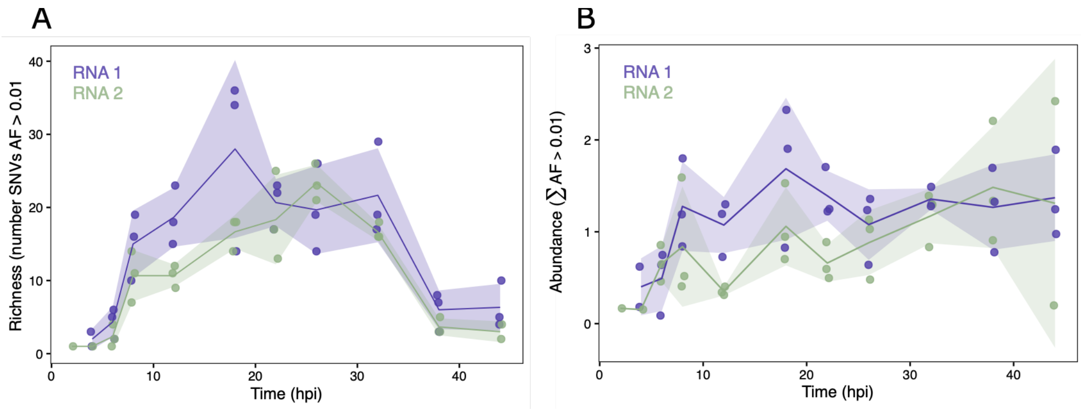


**Figure S2.** SNVs richness and accumulated allele frequency (AF) dynamics. (A) SNV richness as the number of SNVs with allele frequency (AF) > 0.01 per sample. (B) SNV abundance as the sum of AF across all SNVs with AF > 0.01. Lines show the mean profile (±SD) and individual points each sample values. RNA1 in purple and RNA2 in green.

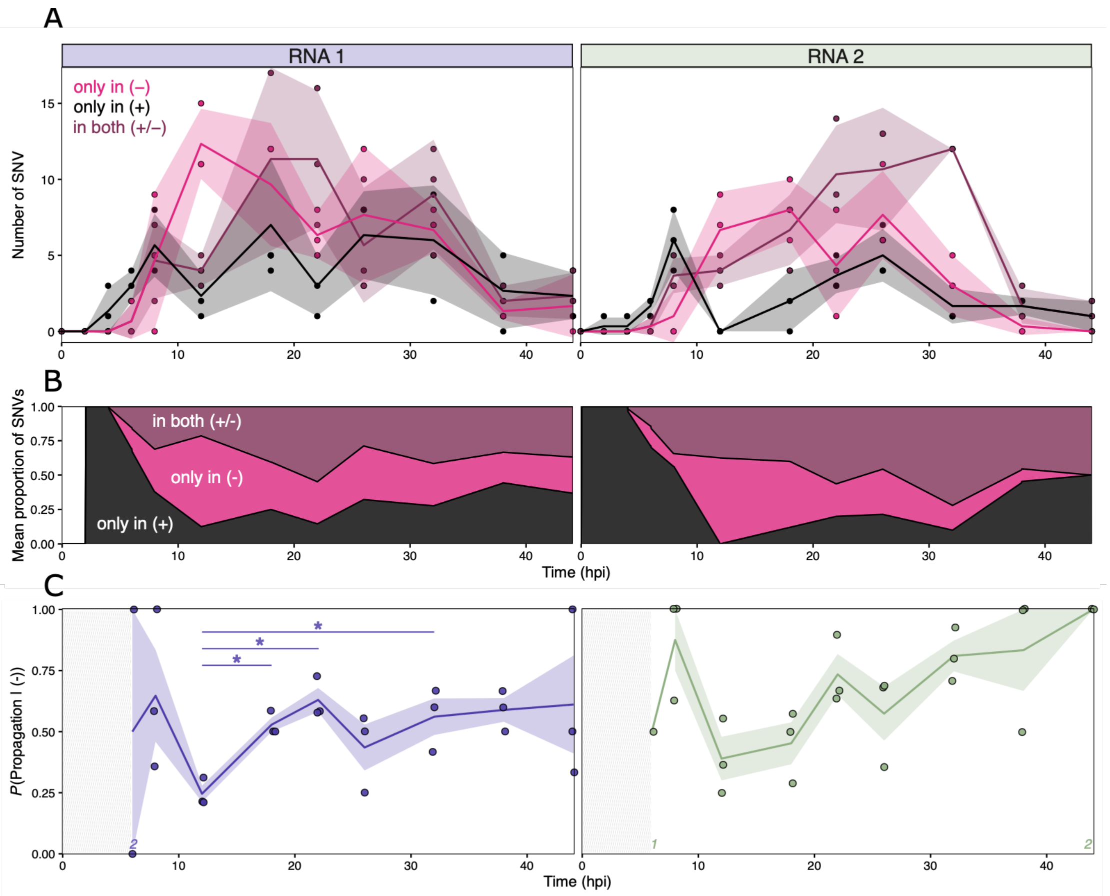


**Figure S3.** Strand-specific SNVs distribution. Composition of SNVs per sample classified as present only in (−)RNA (pink), only in (+)RNA (black) or in both strands (burgundy). **(A)** Number of SNVs (AF > 0.01) of each category in the individual samples. Lines represent mean values (±1 SD). **(B)** Proportion of the mean values per timepoint. **(C)** Propagation probability as the fraction of SNVs detected in both strands over all the SNVs detected in the (−)-strand: $P\left(\text{Propagation}\middle|(-)\right) = \text{both}(+/-)/[\text{both}(+/-) + \text{only}(-)]$. Points represent individual biological replicates; lines indicate the mean transmission probability at each time point and shaded areas represent ±1 SD. Asterisks indicate significant pairwise contrasts between time points derived from estimated marginal means of a quasibinomial generalized linear model with logit link. *P*-values were adjusted for multiple testing using the Benjamini-Hochberg false discovery rate correction (adjusted $P < 0.05$). The number of replicates is indicated in the bottom when they are less than three per timepoint.

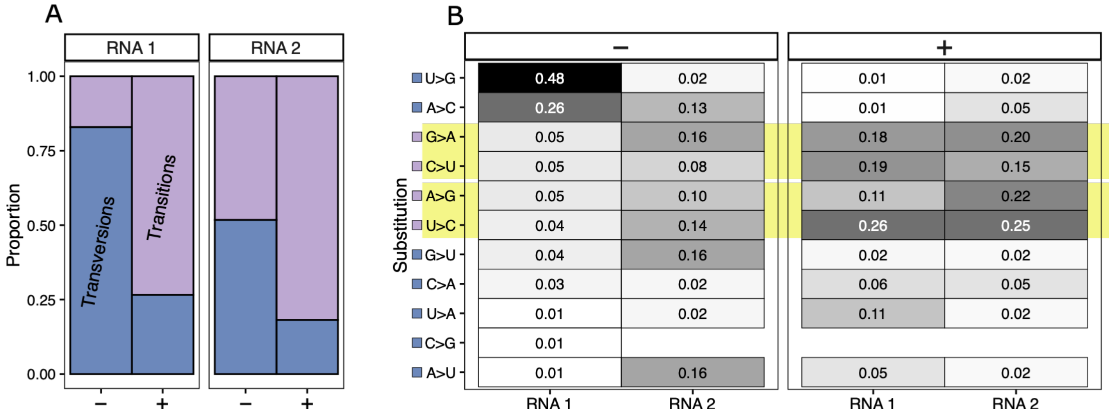


**Figure S4.** Substitution spectrum of strand-specific SNVs. **(A)** Proportional distribution of transitions (purple) and transversion (blue) mutations across strand-specific SNVs. Values are shown separately for (−)RNA- and (+)RNA-associated variants. **(B)** Heatmap of the frequency of individual nucleotide substitutions across RNA segments and strand-specific categories. Each cell shows the proportion of each substitution type within (−)RNA- and (+)RNA-associated SNVs. Putative RNA editing-associated substitutions A>G/U>C (ADAR-like) and C>U/G>A (APOBEC-like changes) are highlighted in yellow for reference. No consistent strand-specific enrichment or pattern indicative of a dominant enzymatic editing signature is observed in the (−)-strand.

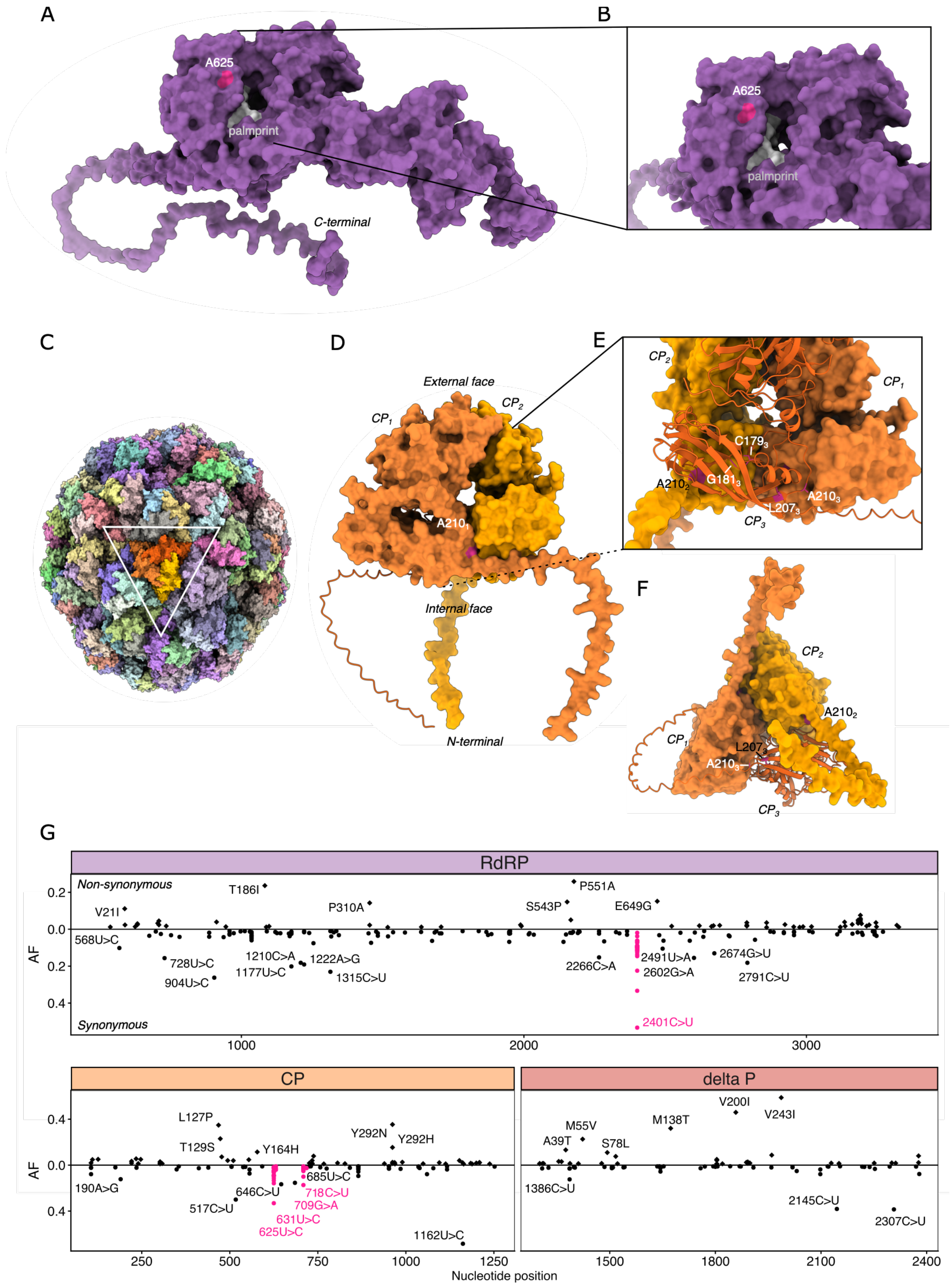


**Figure S5.** Local patterns of $d_N - d_S < 0$ in OrV proteins. **(A)** and **(B)** A 4-codon sliding-window analysis of the $d_N - d_S$ values reveal three hotspots where $d_N - d_S < 0$, one in the RdRP (and two in the CP, corresponding to one and four synonymous SNVs respectively. OrV RdRP was predicted using AlphaFold 3.0. In magenta the residue A625 (correspond to RNA1:2401C>U, in (G). The palmprint conserved motive is represented in light gray. **(C)** OrV viral particle (PBD id: 4NWV). **(D)** - **(F)** The CP trimer was predicted using AlphaFold

3.0 because the N-terminal region is absent from the experimental structure (PBD id: 4NWV). Structural alignment with the experimental model showed close agreement (RMSD = 0.78 Å for 231 pruned atom pairs; 2.80 Å across all 355 pairs). **(G)** Mutational spectrum of all the SNVs used in the analysis with the ones in the selected hotspots in magenta. Only the SNVs founded in the (+) strand with an AF > 0.01 were used. Nonsynonymous SNV are shown in the positive y-axis and synonymous in the negative y-axis.

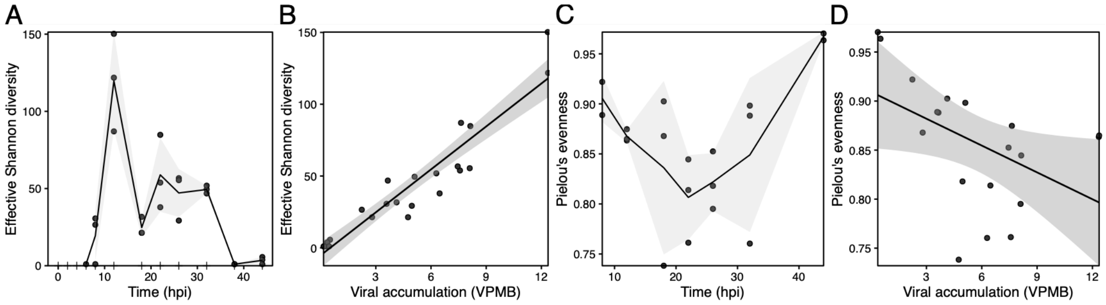


**Figure S6.** Alpha diversity of nsVG communities. **(A)** Temporal dynamics of effective Shannon diversity, expressed as the effective number of equally abundant nsVG clusters. **(B)** Relationship between effective Shannon diversity and viral accumulation (VPMB). **(C)** Temporal dynamics of Pielou's evenness. **(D)** Relationship between Pielou's evenness and viral accumulation. Points represent individual biological replicates. Lines indicate mean values across sampling times (A, C) or linear-model fits (B, D), and shaded areas represent ±1 SD (A, C) or 95% confidence intervals (B, D). Effective Shannon diversity increased strongly with viral accumulation, whereas community evenness showed a slight decline, indicating increasing dominance of a subset of nsVG clusters as viral accumulation increased.

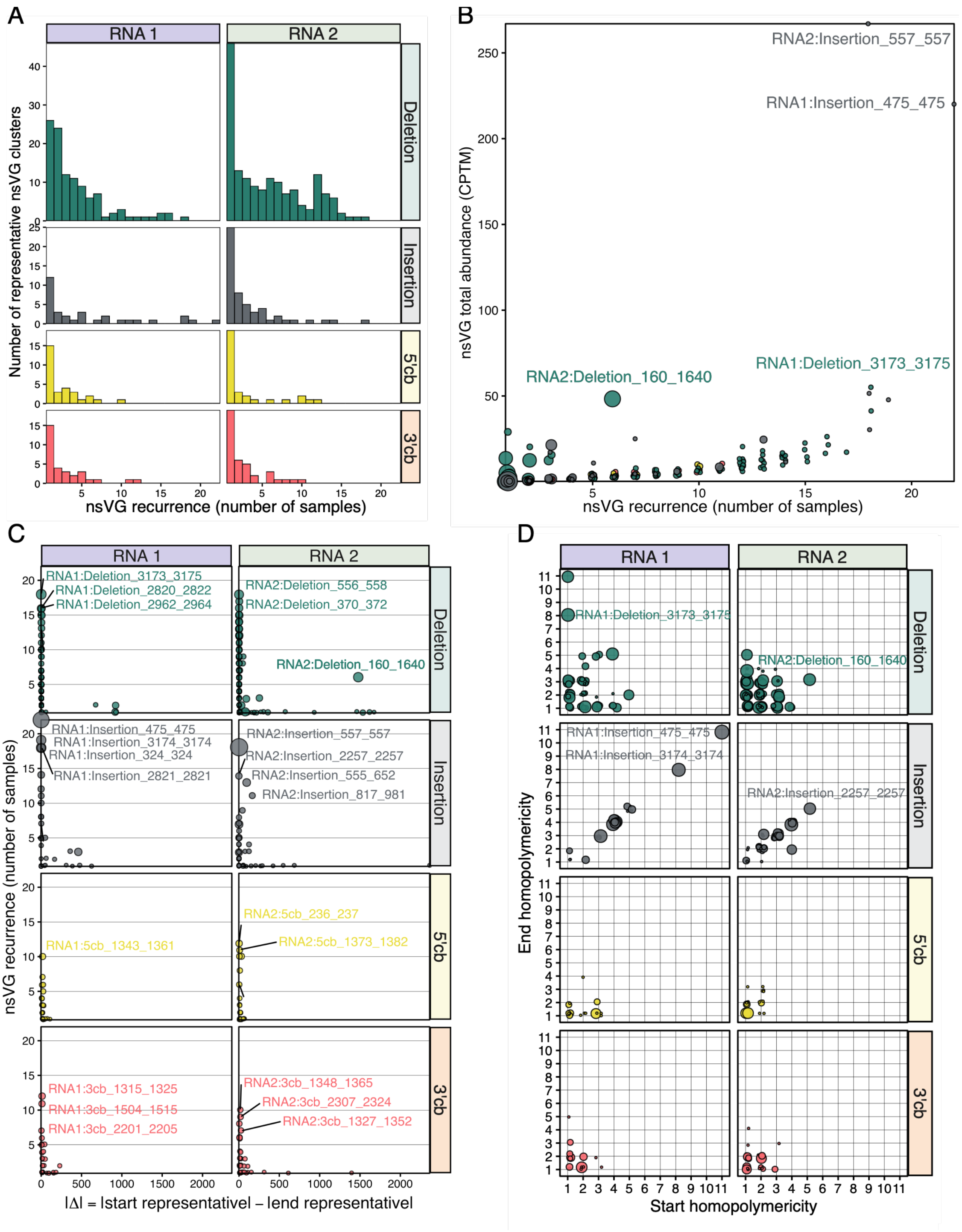


**Figure S7**. Recurrence patterns and sequence context of representative nsVG clusters. **(A)** Distribution of representative cluster recurrence, measured as the number of samples in which each cluster was detected, separated by genomic segment and nsVG category. Most clusters were detected in only a few samples, whereas a limited subset recurred repeatedly across independent infections. **(B)** Relationship between cluster recurrence and total abundance across the dataset. Point size is proportional to the length of the event (|Δ| = |representative end position – representative start position|). Labels indicate highly abundant recurrent clusters. Recurrence was strongly associated with total abundance, showing that a small subset of recurrent clusters contributed disproportionately to the overall nsVG population. **(C)** Relationship between cluster recurrence and event size (Δ). Point size is proportional to total cluster abundance (CPTM). Across all nsVG categories, recurrence decreased with event length ($r_S$ = –0.374, 485 d.f., $P < 0.001$), indicating that highly recurrent nsVGs generally correspond to short genomic modifications, although a small number of large recurrent deletion events were also observed. **(D)** Homopolymeric context of representative nsVG clusters. The homopolymericity of the representative start and end coordinates is shown for each cluster, with point size proportional to recurrence. Recurrent insertion and deletion clusters frequently occurred in or adjacent to homopolymeric tracts.

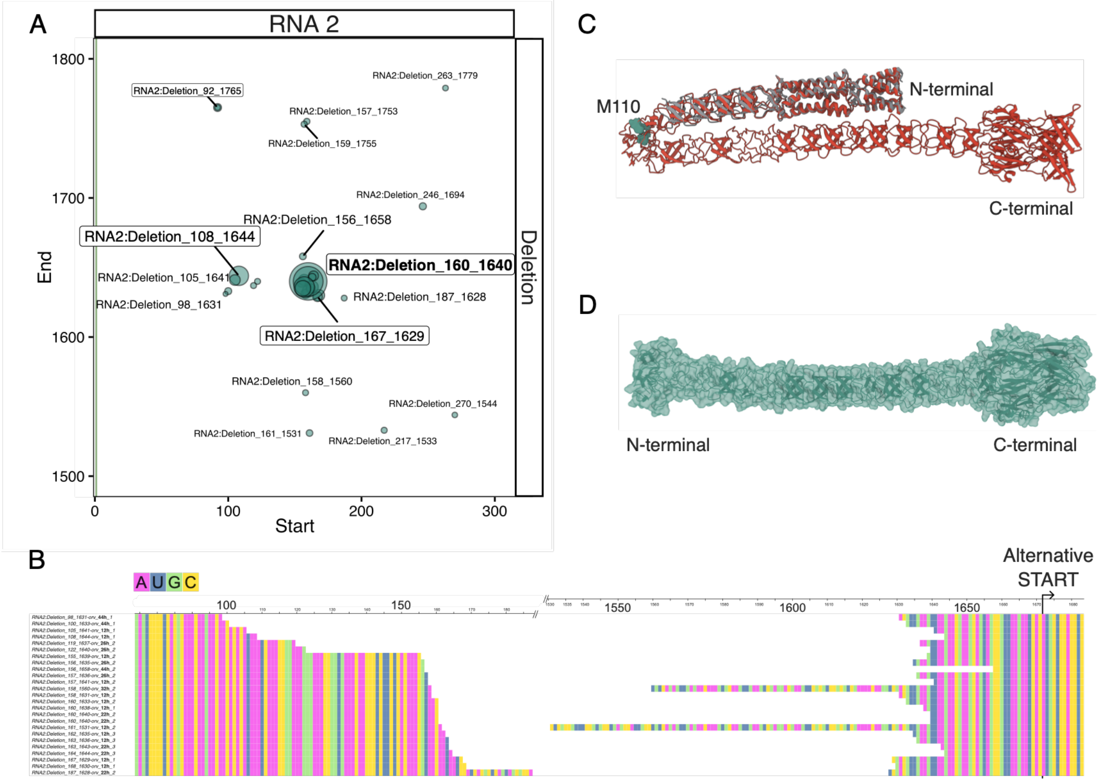


**Figure S8.** Junction heterogeneity within the recurrent RNA2 deletion hotspot and structural prediction. **(A)** Original deletion junctions mapping to the recurrent RNA2 hotspot. Points correspond to individual ViReMa-detected deletion junctions before representative clustering and are plotted using their original start and end coordinates. Point size indicates per sample CPTM abundance. Labels with border mark deletion junctions used as representative cluster IDs, the RNA2:Deletion_160_1640 is also bolded. **(B)** Multiple-sequence alignment of reconstructed deletion-derived RNA2 molecules from the hotspot. Sequences were reconstructed by joining the region upstream of each deletion start to the region downstream of each deletion end. The alignment shows that the hotspot consists of a set of related but non-identical junctions rather than a single fixed deletion. The internal AUG located downstream of the junction region is highlighted; its retention is compatible with the potential expression of a truncated δ product. **(C)** Structural alignment between the experimentally resolved OrV δ N-terminal fragment, residues 1-101 (PDB 5W82) shown in gray, and the AlphaFold 3.0 prediction of the full-length free δ protein, shown in red. The green M110 marks the residue corresponding to the putative internal translation start retained in the deletion-derived RNA2 molecules. **(D)** AlphaFold 3.0 prediction of the protein potentially encoded by the candidate δ sgRNA/nsVG. The model is shown in side view, illustrating the predicted N-terminal globular domain, fiber-like region, and C-terminal domain.

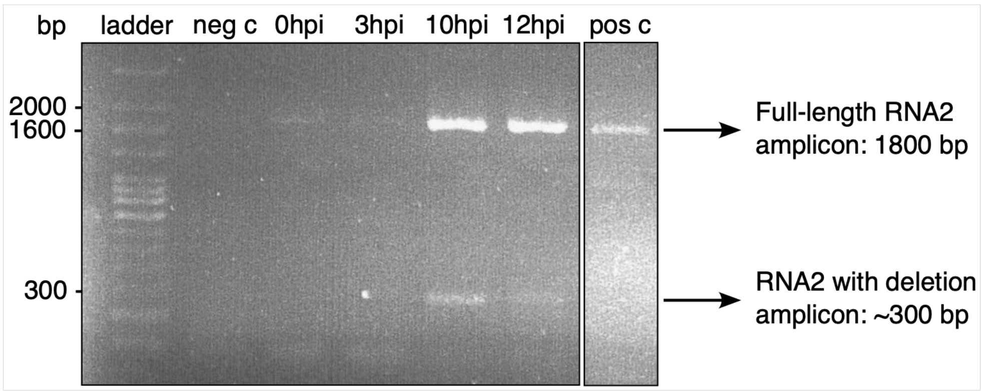


**Figure S9**. Gel electrophoresis of the recurrent RNA2 deletion. RNA2 was amplified by RT-PCR from total RNA of inoculated nematode populations collected at 4 time points post-inoculation (hpi). Primers were designed to span the recurrent deletion (RNA2:Deletion_160_1640) and produce full length (1800 bp) and deletion (~300 bp) amplicons. Total RNA from a healthy worm population was used for the negative control and a plasmid containing full length RNA2 was used for the positive control.
